# Eukaryotic Translation Initiation Factor Loaded Extracellular Vesicles Promotes Macrophage Cholesterol Metabolism in ovarian cancer

**DOI:** 10.1101/2024.12.06.627180

**Authors:** Sonam Mittal, Minal Nenwani, Ishaque Pulikkal Kadamberi, Sudhir Kumar, Olamide Animasahun, Jasmine George, Shirng-Wern Tsaih, Prachi Gupta, Mona Singh, Anjali Geethadevi, Chandrima Dey, Noah Meurs, Ajay Shankaran, Pradeep Chaluvally Raghavan, Deepak Nagrath, Sunila Pradeep

## Abstract

Tumor-driven immune suppression poses a significant impediment to the success of immunotherapy in ovarian cancer. Among the various mechanisms contributing to immune suppression, intracellular communication facilitated by tumor-derived extracellular vesicles (EVs) within the tumor microenvironment (TME) emerges as a pivotal factor influencing tumor growth.

We discovered that EVs from both ovarian tumor cell lines and the plasma of ovarian cancer patients are encapsulated with eukaryotic translation initiation factor 4E (eIF4E). Our study revealed a new mechanism showing how these EVs are loaded with eIF4E and its impact on ovarian cancer progression. We also demonstrated that eIF4E-containing EVs (eIF4E-EVs) alter protein translation in macrophages, contributing to anti-tumor immune response.

Treatment of macrophages with eIF4E-EVs induces an immunosuppressive phenotype marked by the release of cytokines such as IL-6 and an elevated expression of Programmed death-ligand 1 (PD-L1). Notably, eIF4E-packaged EVs enhance the expression of 3-hydroxy-3-methyl-glutaryl-coenzyme A reductase (HMGCR) a pivotal enzyme in cholesterol biosynthesis, resulting in increased cholesterol levels within macrophages. Inhibition of HMGCR or reduction of cholesterol in macrophages effectively restores their antitumor activity by decreasing PD-L1 on macrophages.

Analysis of tumor tissue from ovarian cancer patients revealed a positive correlation between HMGCR and TAM in ovarian cancer. In summary, we have characterized the mechanism of how eIF4E loaded EVs induced cholesterol synthesis, creating an immunosuppressive environment by upregulating PD-L1 expression in macrophages.

## Introduction

Effective communication between cancer cells and surrounding cells within the tumor microenvironment (TME) is a critical factor influencing tumor progression. There has been a growing emphasis on understanding the role of tumor-derived EVs in orchestrating the intricate intracellular communications within the TME ^1-3^. However, the mechanism of EVs crosstalk in TME for oncogenic actions are not well understood. EVs, ranging from 50-300 nm, carry multiple specific proteins or RNAs secreted by host cells^4^. These molecules precisely target recipient cells, influencing tumor behavior and angiogenesis ^5, 6^. Our recent work has reported that tumor-derived EVs cause T-cell exhaustion in TME through sphingosine mediated signaling and impacting immunotherapy outcomes in ovarian cancer ^2^. Emerging evidence suggests that the molecular information contained in EVs can be harnessed for diagnostic and therapeutic purposes^7-11^.

The levels of the individual components of the eIF4F complex, and its activity, are elevated in many human cancers, constituting vulnerability for transformed cells ^12^. The initial recognition of the mRNA 5′-end involves the translation factor eIF4F, a hetero-trimeric complex formed by the assembly of eIF4A, eIF4E, and eIF4G. eIF4E is the cap-binding protein that specifically binds to the 5′-cap structure. eIF4E also interacts with the scaffold protein eIF4G. eIF4G in turn binds to eIF4A, a DEAD-box helicase that has been implicated in the unwinding of double-stranded regions in the 5′-UTR that might otherwise interfere with PIC recruitment and scanning^13^. Early studies demonstrated that eIF4E overexpression enhances the translation of mRNAs with structural repeats in the 5′ untranslated region (5′UTR) in vivo. The average well-translated mRNA has a 5′UTR of 20–50 nucleotides. Around 10% of cellular mRNAs contain atypically long 5′UTR and many of these encode proto-oncogenes, anti-apoptotic proteins, and growth factors. A long 5′UTR and GC rich sequence tends to form a stable secondary structure. It has been shown that translation of mRNA with long 5′UTR is often sensitive to expression level of eIF4E and eIF4A1.While it is established that endogenous levels of eIF4F complex proteins are elevated in the TME, there is limited reporting on their encapsulation within EVs. Our data shows that the presence of eIF4E in ovarian cancer derived EVs, underscoring its significance in these vesicles. This mechanism of eIF4E encapsulation in EVs holds significant implications as it can facilitate the delivery of eIF4E into other cells in the TME, thereby augmenting tumorigenic potential.

The TME comprises a dynamic cellular environment surrounding the tumors, encompassing macrophages, stroma, stem cells, fibroblasts, lymphocytes, pericytes, adipocytes, and blood vessels ^14^. Tumor-associated macrophages (TAMs) represent the majority of immune cells found in ascites, making them the predominant immune cell type in ovarian cancer ^15^. Tumor cell released EVs influence macrophages, modulating immunity, regulating inflammation, and impacting the TME ^16^. Therefore, understanding macrophage dynamics in ovarian cancer ascites is crucial for developing targeted therapeutic approaches to disrupt the tumor-promoting environment and improve treatment outcomes.

Our data demonstrate that tumor derived EVs, through the action of eIF4E, impact the protein synthesis in macrophages, notably augmenting the translation of genes associated with cellular metabolism. Furthermore, our findings elucidate that eIF4E-EVs promote tumor formation by inducing the accumulation of cholesterol in macrophages via 3-hydroxy-3-methyl-glutaryl-coenzyme A reductase (HMGCR), resulting in an immunosuppressive phenotype in the TME. Therefore, inhibiting HMGCR in macrophages or reducing cholesterol levels within the TME will have the potential to suppress tumor growth. Our research sheds light on the mechanism whereby tumor cells exploit EVs to transport eIF4E, a critical translation-regulating molecule, to macrophages, thereby dampening anti-tumor immune responses.

## Results

### 1. eIF4E is transported by ovarian tumor derived EVs

Numerous studies have shown that ovarian cancer cells release a significant number of EVs into the patients’ ascites and plasma. These EVs, specifically derived from tumor cells are anticipated to carry a molecular signature that partly mirrors that of the parental tumor cells ^17^. Furthermore, these EVs are enriched in molecules known to promote tumor growth ^17, 18^. We conducted a comprehensive proteomic analysis to understand the impact of EVs derived from ovarian cancer cells on tumor growth and metastasis. We isolated EVs from OVCAR5 cells and characterized them using nano tracking analysis and transmission electron microscopy (TEM) which confirmed them within the size range expectations (50-300 nm). Their morphology is consistent with the standard established in MISEV (Figures S1A and 1B). Additionally, the presence of EVs in our preparations was validated by the detection of proteins typically enriched in EVs (ALIX, TSG101, CD63) and the exclusion of intracellular proteins that are not shed in EVs (GM130) by western blot (Figure S1C).

Our mass spectrometric analysis, using liquid chromatography-tandem mass spectrometry (LC-MS/MS) on EVs isolated from HeyA8 and OVCAR5 cells, identified a total of 2,249 proteins common to the EVs from both cell lines (Figures1A, 1B and Table S1). The results from Ingenuity Pathway Analysis (IPA) underscored proteins that were highly enriched in pathways related to eukaryotic translational regulation (Figure 1C).

**Figure 1:**
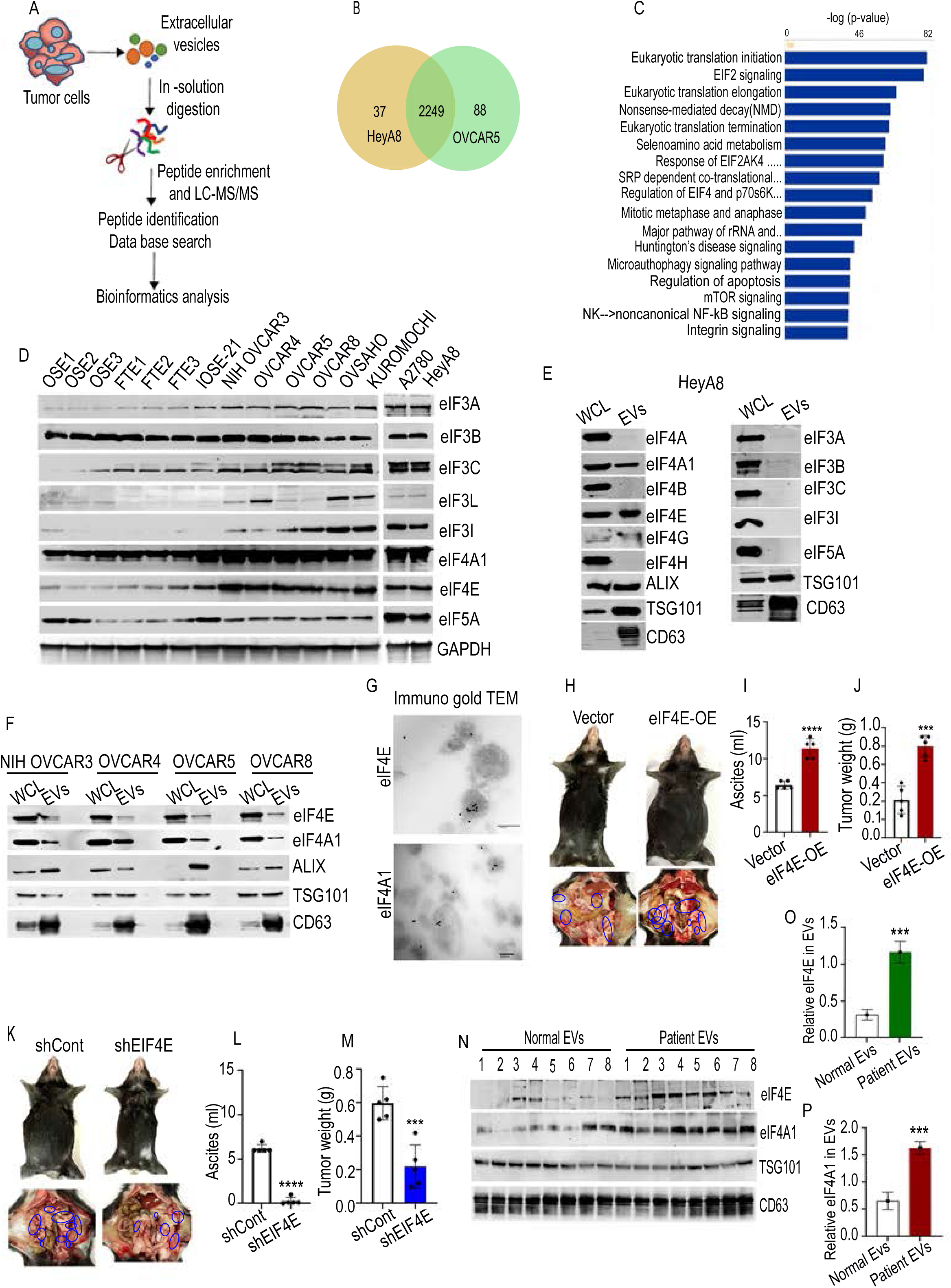
Extracellular vesicle isolation and proteomic analysis. (A) EV isolation and proteomics workflow. (B) Venn diagram of total proteins identified in HeyA8 and OVCAR5 cell-derived EVs by LC-MS/MS. (C) Pathway analysis for proteins identified in HeyA8 and OVCAR5 cell derived EVs using Ingenuity Pathway Analysis (IPA). The top pathways were ranked based on p-value, and the bars represent the inverse log of the p-value (x-axis). (D) Expression of most abundant eukaryotic translation initiation proteins identified. (E and F) Western blot analysis for different eIFs proteins on isolated EVs and whole cell lysates. (G) Representative TEM image of OVCAR8 cells derived EVs. EVs were immunogold-labeled with anti-eIF4E and anti-eIF4A1 antibodies. Scale bar, 200nm and 100nm. (H) Enlarged peritoneal cavity and increased tumor burden (blue circles) in eIF4E overexpressed ID8Trp53^-/-^;Brca2^-/-^ cells. n=5 mice/group. (I) ascites volume and (J) tumor weight. (K) Representative image of the mouse from sh-control and sh-EIF4E group. Peritoneal cavity of mice showing tumor locations (blue circles). n=5 mice/group. (L) ascites volume and (M) tumor weight. (N) Validation of different proteins in EVs isolated from plasma of healthy donors and ovarian cancer patients. (O) Densitometry analysis of experiment as in Figure N. Error bars indicate mean ± SEM. Significance was determined by Student’s t test, where ∗∗∗p < 0.001, ∗∗∗∗p < 0.0001.

To narrow down our search for a specific translational regulator, we evaluated the protein expression within these pathways across various ovarian cancer cell types. Among the translational regulators enriched in EVs from HeyA8 and OVCAR5 cells, the eIF3, eIF4, and eIF5A proteins stood out as the most abundant. Immunoblotting confirmed that these proteins were higher in ovarian cancer cells compared to primary ovarian surface epithelial (OSE) and fallopian tube surface epithelial (FTE) cells (Figure 1D). We sought to determine the presence of “eIF3”, “eIF5A” and “eIF4” factors in EVs isolated from HeyA8 cells to validate the mass spectrometry data. Immunoblotting confirmed the presence of eIF4E and eIF4A1 in both whole cell lysate and EVs isolated from the supernatants of HeyA8 cells. However very less expression of eIF4G was reported in EVs from HeyA8 cells. (Figure 1E). eIF4E and eIF4A1 were also present in EVs isolated from multiple human and murine ovarian cancer cell lines (Figures 1F and S1D). However, eIF4E and eIF4A1 were minimally expressed in EVs isolated from OSE and IOSE-21 cells (Figures S1E and S1F). TEM images using immunogold staining for eIF4E and eIF4A1 confirmed the presence of these proteins within tumor cells derived EVs (Figure 1G).

To ensure the purity of the EV preparations enriched in eIF4E and eIF4A, we employed sucrose gradient centrifugation, a widely recognized EVs fractionation technique that separates EVs from soluble proteins and nucleic acids^19^. In line with established EV fractionation protocols, fractions 3-5 (of 8), representing the 20-40% sucrose gradient, exhibited the highest levels of EV markers (CD63 and ALIX), along with eIF4E and eIF4A1 (Figure S1G), consequently, these fractions were selected for all our future experiments.

To confirm the notion that eIF4E and eIF4A1 are indeed transported by EVs, we selectively inhibited EVs release from tumor cells by using the EVs biogenesis inhibitors dimethyl amiloride (DMA) (20 µg/ml)^20^. DMA significantly reduced the level of eIF4E and eIF4A1 in EVs, suggesting that blocking EVs biogenesis inhibits the packaging of these proteins inside EVs without affecting the intracellular levels. (Figures S1H and S1I).

To elucidate the distinct functions of eIF4E and eIFA1, we altered their expression levels within cells to generate EVs enriched or depleted in these proteins, alongside other EV constituents. Our *in vivo* data demonstrated that C57BL/6 female mice implanted with ID8Trp53^-/-^;Brca2^-/-^ murine ovarian cancer cell lines overexpressing eIF4E or eIF4A1 exhibited an increased volume of ascites and tumor burden within the peritoneal cavity compared to the control group (Figure 1H-1J, and S1J-M). Similarly, mice implanted with eIF4E and eIF4A1 knockdown (KD) ID8Trp53^-/-^;Brca2^-/-^ cells (Figures S1N-S1O) had a decreased volume of ascites and tumor burden as compared to the control group (Figures 1K-1M, and S1P-1R). Our IHC and western blotting analysis also revealed a significant increase in eIF4E and Ki67 levels in tumor tissue after eIF4E overexpression (Figure S1S) and vice versa in eIF4E KD group (Figure S1T).

Moreover, we observed the presence of eIF4E and eIF4A1 in EVs isolated from the plasma of ovarian cancer patients, contrasting with their minimal expression in EVs from healthy donors (Figure 1N-1P). Additionally, we confirmed the presence of EV proteins TSG101, and CD63 in vesicles obtained from both ovarian cancer cells and patient plasma.

### 2. EVs packaged with translation initiation factors enhance global protein synthesis

Proteins associated with the endosomal sorting complex required for transport (ESCRT), such as TSG101, are responsible for recruiting cargo proteins into EVs ^21^. We used a GFP-trap co-immunoprecipitation assay to understand whether eIF4E and eIF4A1 are packaged into EVs by directly interacting with ESCRT proteins. GFP-tagged beads were employed to precipitate eIF4E-GFP and eIF4A1-GFP from the lysates of OVCAR8 cells, which stably express eIF4E and eIF4A1 GFP fusion proteins (Figure 2A). This assay revealed a unique interaction between the ESCRT-I component, TSG101 and both eIF4E and eIF4A1. To confirm these interactions, reciprocal immunoprecipitation was performed using OVCAR8 cells expressing TSG101-GFP. The results showed that TSG101 co-immunoprecipitated with eIF4A1 and eIF4E, confirming its interaction with these proteins and their encapsulation into EVs (Figure 2B).

**Figure 2.**
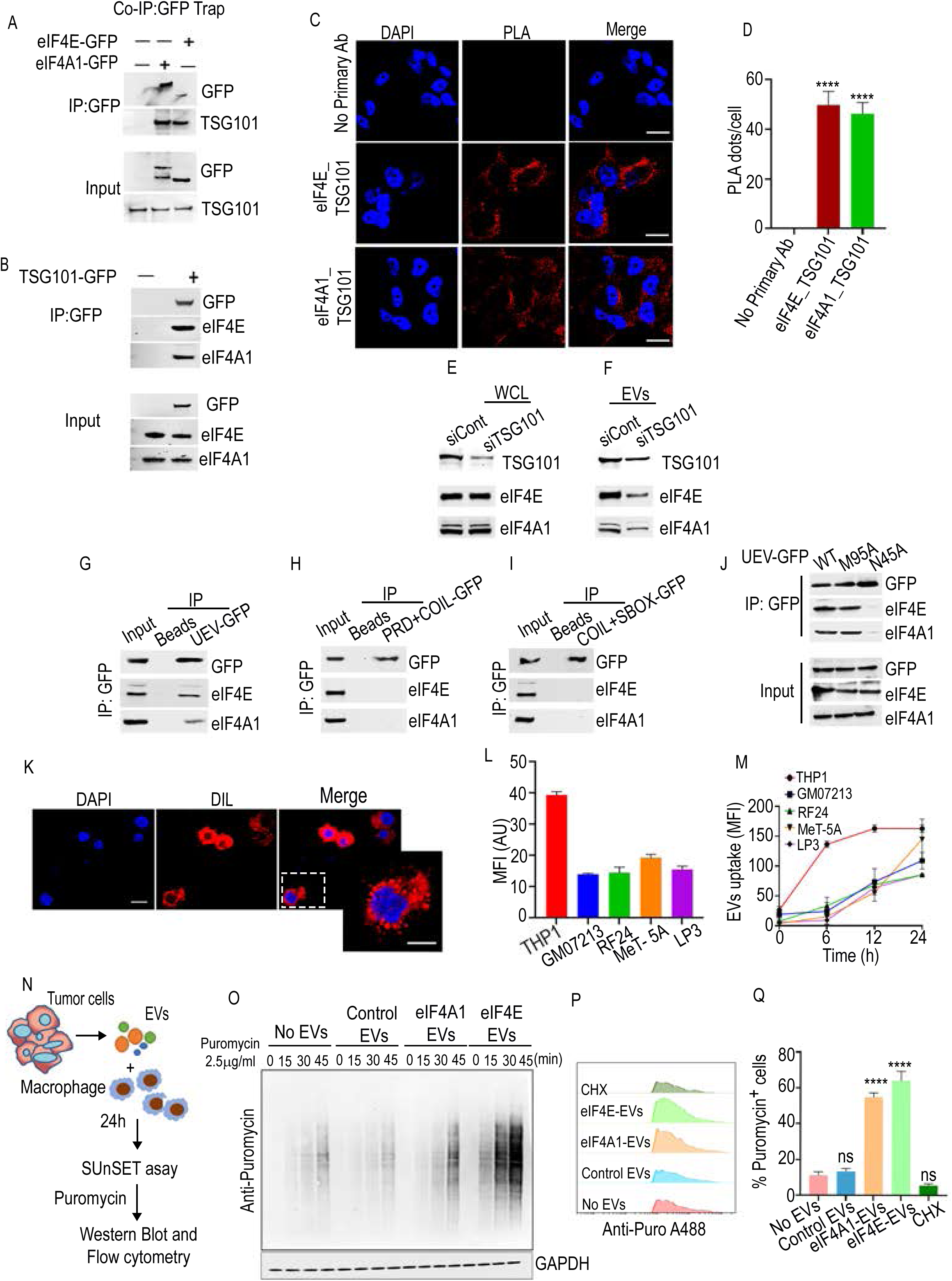
Translation initiation factors in EVs enhance protein synthesis in macrophages. (A) eIF4E-GFP, eIF4A1-GFP fusion proteins and their binding partner TSG101 was immunoprecipitated with GFP-tag beads. (B) Reverse immunoprecipitation of TSG101-GFP fusion protein and its interacting partners, eIF4E and eIF4A1. (C) In-situ proximity ligation assay (PLA). Red dots represent the interaction of eIF4E and eIF4A1 with TSG101. Nuclei were stained with DAPI (blue). Scale bar 50µm. (D) Bar graph represents the mean of red PLA dots with SEM counted using image J software. (E and F) Immunoblot analysis of the WCL and EVs fraction after TSG101 knockdown in OVCAR8 cells. (G) Co-IP showing of TSG101-UEV-GFP, eIF4E and eIF4A1. (H) Co-IP of TSG101-PRD+COIL-GFP, eIF4E and eIF4A1. (I) Co-IP of TSG101-COIL+SBOX-GFP, eIF4E and eIF4A1. (J) Western blotting of Co-IP assay of TSG101-UEV wildtype (WT) and mutants (M95A, N45A) with eIF4E and eF4A1. (K and L) DIL-labeled EVs uptake in different cells. Scale bar 50µm and 20µm. (M) DIL-labeled EVs uptake kinetics. (N) Schema illustrates the principle of surface sensing of translation (SUnSET) assay. (O) SUnSET measurements of protein synthesis by western blotting in THP1-derived macrophages. (P and Q) FACS of the puromycin-labeled cells using the anti-puromycin antibody tagged with Alexa Fluor 488. Error bars indicate mean ± SEM, Significance was determined by Student’s t test and one-way ANOVA where ****p<0.0001. ns, non-significant.

To further investigate the interaction between eIF4E and eIF4A1 with TSG101, we employed proximity ligation assay (PLA) to detect proximity between eIF4E-TSG101and eIF4A1-TSG101. There was a significant increase in the PLA intensity when primary antibodies against the two binding partners, either eIF4E or eIF4A1 and TSG101, were used, indicating the physical closeness between both eIF4E-TSG101 and eIF4A1-TSG101 (Figures 2C and 2D).

Next, we sought to determine whether TSG101 plays a role in recruiting eIF4E and eIF4A1 into EVs. To accomplish this, we depleted TSG101 by siRNA in OVCAR8 cells. Supernatant was collected and analyzed for eIF4E and eIF4A1 in EVs. Western blotting revealed a significant reduction in the levels of eIF4E and eIF4A1 in EVs isolated from TSG101 knockdown cells but not in the whole cell lysate, indicating the involvement of TSG101 in recruiting eIF4E and eIF4A1 into EVs (Figures 2E and 2F).

Our subsequent objective was to identify the TSG101 domain essential for its binding to eIF4E and eIF4A1, thereby facilitating their sorting into EVs. Previous studies have established that the ubiquitin-E2-like variant (UEV) domain of TSG101 protein interacts with the ubiquitin moiety of proteins designated for sorting or degradation ^22-24^. For instance, TSG101 recognizes ubiquitin through the UEV domain, which is required for sorting EGFR into multivesicular bodies (MVBs) ^23, 25^. To address this, we conducted Co-IPs with three truncated TSG101 proteins (Figures S2A-S2C). Our results identified that only UEV domain directly interacts with eIF4E and eIF4A1 (Figures 2G-2I). Further experiments showed that mutations in TSG101 UEV domain amino acids, known to be crucial for TSG101 interaction with ubiquitin (N45A), abolished the interaction of TSG101 with eIF4E and eIF4A1, whereas the replacement of other amino acids in the UEV domain (M95A) of TSG101 had no distinct effect (Figure 2J). Taken together, our data provide compelling evidence that the UEV domain is important for the binding of TSG101 to eIF4E and eIF4A1, thus facilitating their encapsulation into EVs.

Following this, we determined the uptake efficacy of tumor cells derived EVs by various cell types. The EVs were isolated from OVCAR8 cells and fluorescently labeled with DIL dye. These labeled EVs were subsequently incubated with THP1-derived macrophages, fibroblasts (GM07213), endothelial (RF24), and mesothelial (MeT5A, LP3) cells (Figures 2K-2L and S2D). Our observations revealed that macrophages exhibited increased internalization of EVs compared to the other cell types. Furthermore, we delved into the dynamics of internalization. Consistent with the above observations, the temporal patterns of EV internalization exhibited two distinct patterns. In the case of macrophages, the uptake of EVs significantly increased over time, reaching a plateau after 6 h. Conversely, for other cell types, efficient detection of EV uptake was observed at 24 h of incubation (Figures 2M and S2E).

To elucidate the impact of EV-packaged eIF4E/eIF4A1 on global translational regulation, macrophages, endothelial cells, and fibroblast cells were incubated for 24 h with EVs isolated from (I) parental OVCAR8 and OVCAR5 cells, (II) OVCAR8 and OVCAR5 cells that overexpress eIF4A1 or eIF4E). We assessed total protein synthesis using the Surface Sensing of Translation (SUnSET) assay, which employs an anti-puromycin antibody for immunological detection ^23^. During a newly synthesized polypeptide chain, puromycin is incorporated into the nascent polypeptide chain, thereby inhibiting the formation of a new peptide bond with the subsequent aminoacyl-tRNA and terminating peptide elongation (Figure 2N). This leads to the release of a truncated puromycin-bound peptide from the ribosome. We performed two separate experiments utilizing the SUnSET assay principle. We performed immunoblotting with puromycin antibody and immunostaining with fluorochrome-labeled anti-puromycin antibody, followed by flow cytometry. Both the assays demonstrated an increase in protein synthesis rates, notably in macrophages compared to the other cell types (Figures 2O-2Q and S2F-S2M). These findings prompted us to hypothesize that macrophages undergo significant reprogramming when exposed to EVs released by ovarian cancer cells within the tumor microenvironment.

### 3. eIF4E-loaded EVs induce metabolic reprogramming in macrophages

To gain a comprehensive understanding of the most affected biological processes and the proteins that are either upregulated or downregulated by eIF4A1 and eIF4E loaded EVs, we performed SILAC proteomic analysis in cultured THP1-derived macrophages (Figure 3A). We treated macrophages with EVs derived from OVCAR8 cells and performed mass spectrometry. In this analysis, we identified a total of 1639 proteins using the SILAC-MS in THP1-derived macrophages. Among these, 235 proteins exhibited significant upregulation, while 458 proteins showed significant downregulation in response to eIF4E-EVs compared to control EVs (Table S2). Our Gene ontology (GO) analysis indicated that the pathways related to phagosome maturation and metabolic alterations were specifically enriched in the eIF4E-EV treated group (Figure 3B). However, no metabolically altered genes or pathways were found in the eIF4A1-EV treated group. This could be because availability and/or activity of individual eIF4F component has been shown to have a selective impact on the translation of specific mRNAs. Although the mechanisms underlying this selectivity are incompletely understood, features within the 5′ UTR appear to be a main determinant. For instance, the translation of mRNAs with very short 5′ UTRs is disproportionally reduced, compared with global translation rates, upon inhibition of eIF4E activity, while remaining relatively insensitive to the perturbation of eIF4A activity. Subsequently, we identified the proteins with a ±1.2-fold difference associated with these pathways in our SILAC dataset and generated a heat map. Proteins associated with key metabolic pathways such as glucose (HK2), glutamine (GLS), and cholesterol metabolism (HMGCS1, LIMA1) were upregulated, while proteins involved in phagocytosis and endocytosis (DNM1L, DNM2, AP2A1, and AP3B1) displayed downregulation (Figure 3C).

**Figure 3.**
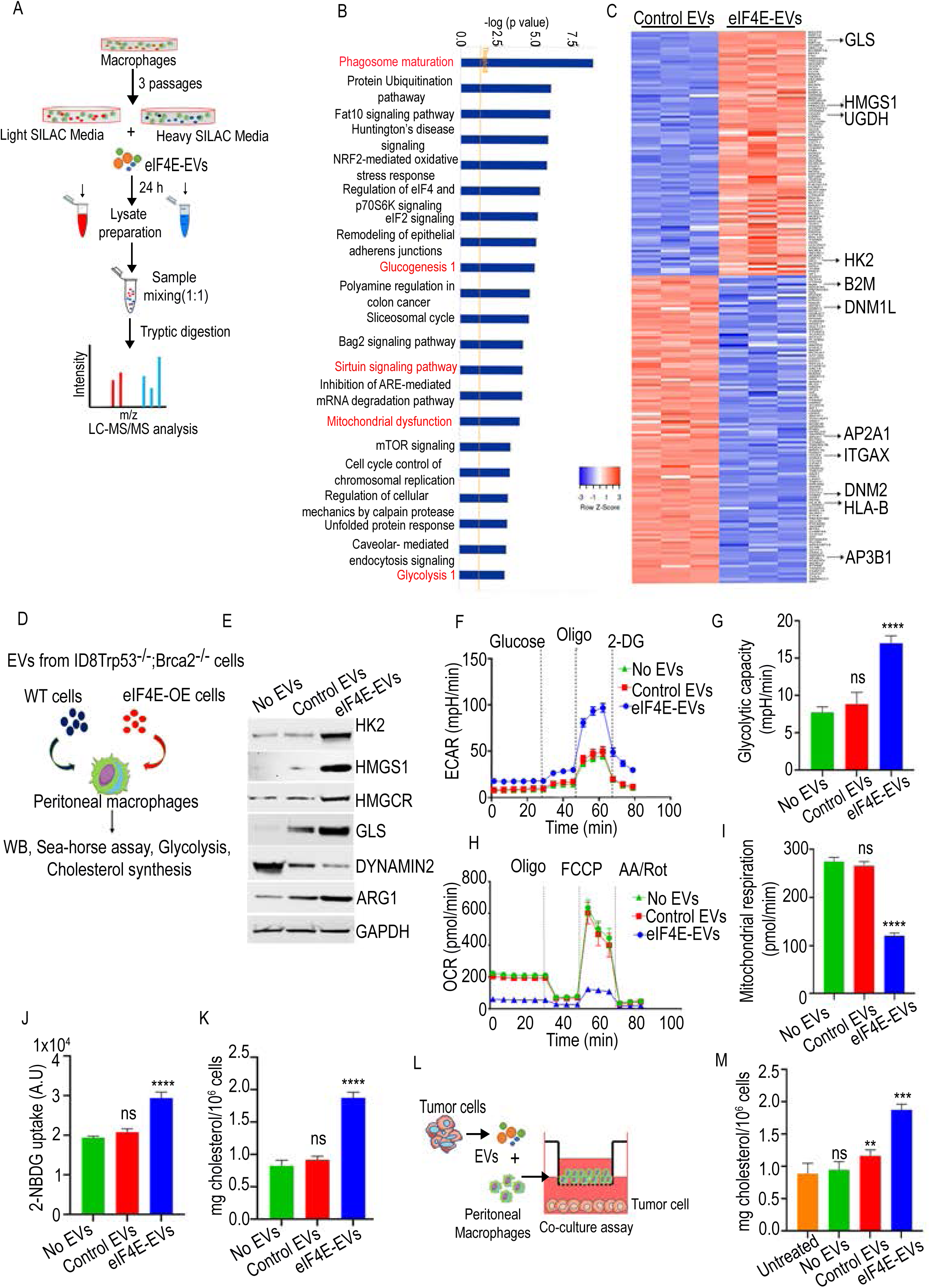
eIF4E-loaded EVs induce metabolic reprogramming in macrophages. (A) Schema representing SILAC-MS workflow. (B) IPA analysis of proteins identified in THP1-derived macrophages treated with EVs. (C) Heat map showing significantly upregulated and downregulated proteins. (D) Workflow representing validation of SILAC data. (E) Representative western blots showing expression of HK2, HMGCS1, HMGCR, GLS, ARG-1, Dynamin-2, in peritoneal macrophages treated with EVs. (F and G) Seahorse glycolysis stress test with sequential addition of glucose, oligomycin, and 2-DG in peritoneal macrophages pre-treated with eIF4E-EVs or control EVs. (H and I) OCR measurement in tumor derived EVs-stimulated macrophages. (J) Glucose uptake assay with 2-NBDG in macrophages stimulated with EVs. (K) Cholesterol synthesis in macrophages by Amplex cholesterol assay. (L) Schema of co-culture assay. (M) Cholesterol estimation in co-culture assay. Data are representative of three independent experiments. Error bars indicate mean ± SEM, *p<0.05, **p<0.01, ***p<0.001, ****p<0.0001 (one-way ANOVA). ns, non-significant.

The SILAC data validation involved western blot analysis on peritoneal macrophages isolated from naïve mice treated with EVs from WT and eIF4E-overexpressing cells. eIF4E-EVs induced increased expression of HK2, GLS, HMGCS1, and HMGCR. Both HMGCS1 and HMGCR are the rate-limiting enzymes in the mevalonate pathway for cholesterol synthesis ^26^. Additionally, eIF4E-EV stimulation elevated ARG-1 levels in macrophages and decreased DNM2 (Dynamin 2) (Figures 3D and 3E). Arg1 is a key driver of immune suppression found predominantly in TAM and Dynamin 2 is required for phagocytosis in macrophages ^27^,^28^. Similar findings were validated in THP1-derived macrophages (Figures S3A and S3B).

Seahorse assays, probing glycolytic and mitochondrial functions demonstrated the metabolic shifts triggered by eIF4E-EVs in macrophages. Intriguingly, peritoneal macrophages treated with eIF4E-EVs showcased heightened extracellular acidification rates (ECAR) relative to the control EV group (Figures 3F and 3G). Additionally, analysis of the oxygen consumption rate (OCR) unveiled a decline in mitochondrial respiration capacity among macrophages treated with eIF4E-EVs (Figures 3H and 3I). These findings were consistently observed in THP1-derived macrophages exposed to both control and eIF4E-EVs (Figures S3C-S3F).

In contrast, treatment with eIF4A1-EVs had no significant effect on ECAR and OCR when compared to control EVs (Figures 3G and 3H). Additionally, we conducted a glucose consumption assay, where glucose-deprived macrophages were co-cultured with 2-NBDG, a glucose analog. Peritoneal and THP1-derived macrophages stimulated by eIF4E-EVs showed a significant increase in glucose uptake compared to those treated with control EVs (Figures 3J and S3I). In addition to glycolysis assays, we quantified cholesterol in the EV treated macrophages which revealed a significant increase in cholesterol levels following treatment with eIF4E-EVs (Figures 3K and S3J). Conversely, macrophages treated with eIF4A1-EVs exhibited no increase in glucose uptake and cholesterol levels compared to those treated with control EVs (Figures S3K and S3L). This finding further corroborates the SILAC analysis, indicating no significant changes in the expression of genes associated with the glucose and cholesterol pathways upon eIF4A1-EVs treatment. Consequently, our attention shifted towards exploring the translational and metabolic alterations specifically induced by eIF4E-EVs in macrophages for subsequent studies.

To elucidate the paracrine signaling interactions between cancer cells and TAM, we co-cultured ID8Trp53^-/-^;Brca2^-/-^ with peritoneal macrophages treated with eIF4E-EVs under various conditions for 48 h. We observed that tumor cells co-cultured with eIF4E-EV treated macrophages exhibited significantly elevated cholesterol levels when compared to both the no EVs and control EV group (Figures 3L and 3M). Similarly, when OVCAR8 cells were co-cultured with THP1-derived macrophages treated with EVs, we observed an increase in cholesterol levels in OVCAR8 cells within the eIF4E-EVs group (Figures S3M and S3N). These collective findings underscore the pivotal role of tumor derived EVs packed with eIF4E in reprogramming macrophage metabolism.

### 4. Tumor derived EVs with eIF4E encapsulation enhance *de novo* cholesterol synthesis in macrophages

We observed that treatment of macrophages with eIF4E-EVs increased the expression of enzymes associated with glucose, glutamine, and cholesterol metabolism (Figure 3E). As protein synthesis directly depends upon the amino acid generation via tricarboxylic acid (TCA) cycle, we wanted to check the incorporation of amino acids and their metabolites as well as the consumption of amino acids toward protein synthesis after EVs treatment. We conducted untargeted metabolic analysis on macrophages exposed to tumor derived EVs. Interestingly, our global metabolic assessments aligned with the upregulated protein synthesis in macrophages treated with eIF4E-EVs. Specifically, we observed elevated levels of essential amino acids - valine, methionine, threonine, tryptophan- and non-essential amino acids -asparagine, glutamine, aspartate, glutamate-which are important for protein synthesis during translation (Figures 4A-4J). The upregulation of protein translation is known to be coupled with nucleotide biosynthesis, facilitated by eIF4E mediated regulation of phosphoribosylpyrophosphate synthetase (PRPS2) ^29^. PRSP2 is a rate limiting enzyme of the pentose phosphate pathway, exerting control over nucleotide biosynthesis. Interestingly, our untargeted analysis unveiled an augmentation in pentose phosphate pathway intermediates, such as sedoheptulose 7-phosphate and 6-phosphogluconate. Additionally, we observed an overall elevation in purines, pyrimidines and intermediates of TCA cycle in macrophages treated with eIF4E-EVs (Figures 4A, 4B and S4A-S4M). We noted intriguing changes in the lipid metabolism of macrophages treated with eIF4E-EVs. Specifically, we observed an increase in immunosuppressive polyunsaturated fatty acids (oleic and linoleic acid) alongside a decline in saturated fatty acid (palmitic acid) levels. Consistent with our SILAC data analyses, we also observed elevated levels of geranyl pyrophosphate, a precursor in the cholesterol synthesis pathway, in eIF4E-EV treated macrophages compared to control macrophages (Figures 4A and 4B). Overall, these systemic metabolic changes, combined with protein expression analysis, hinted at an increase in amino acid, nucleotide, and cholesterol biosynthesis in EVs treated macrophages.

**Figure 4.**
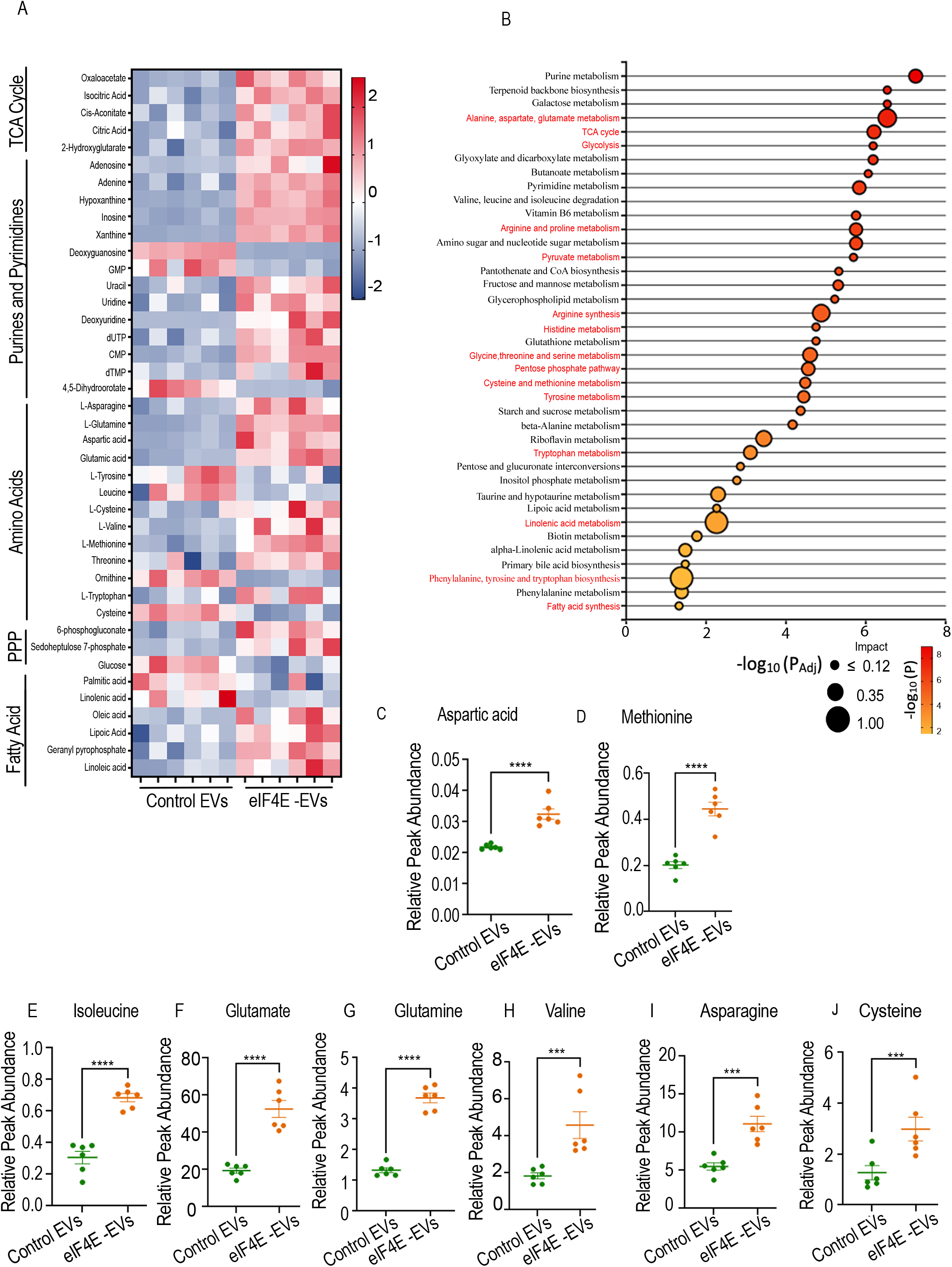
Effect of tumor derived EVs on global metabolomics. (A). Heat-map of untargeted metabolic analysis showing z-scores of differentially upregulated and downregulated metabolites. n=6. (B) Pathway enrichment analysis based on human metabolic pathways in the KEGG database. n=6 (C and J) Relative metabolites levels (peak intensity normalized by internal standard and cell number between control and eIF4E-EVs treated macrophages. n=6. The data are shown as mean ± SEM. ***p<0.001 ****p<0.0001 compared to control by Student’s t-test and one way ANOVA.

Further, to unravel the contribution of glucose or glutamine metabolism towards lipogenesis we carried out semi-targeted stable isotope tracing analysis of EVs treated macrophages using gas chromatography mass spectrometry. ^13^C labelled glucose tracing revealed an increased amount of glycolytic and TCA cycle intermediates synthesized from glucose in EVs treated macrophages. More specifically, we noted elevated levels of M+3 phosphoenolpyruvic acid (PEP) and pyruvate. A similar trend was seen in the TCA metabolites such as malate, fumarate, and citrate (Figure 5A). Furthermore, several other metabolites including alanine, aspartate, 2-hydroxyglutarate, cytosine, and proline are also synthesized in greater amounts from labeled glucose (Figures S5A-S5E). Therefore, it can be inferred that macrophages treated with eIF4E-EVs are actively utilizing the glycolytic pathway for biosynthesis and ATP generation as compared to control EVs. Conversely, glutamine tracing experiments revealed no significant difference in glutamine utilization between the two groups (Figure S5F). To determine the source of the elevated cholesterol levels in treated macrophages (Figure 3K), we utilized ^13^C labelled glucose tracing for 96 h. Remarkably, we observed an increase in *de novo* cholesterol synthesis, as evidenced by the heightened incorporation of labeled glucose into cholesterol in macrophages treated with eIF4E-EVs (Figures 5B and 5C).

**Figure 5.**
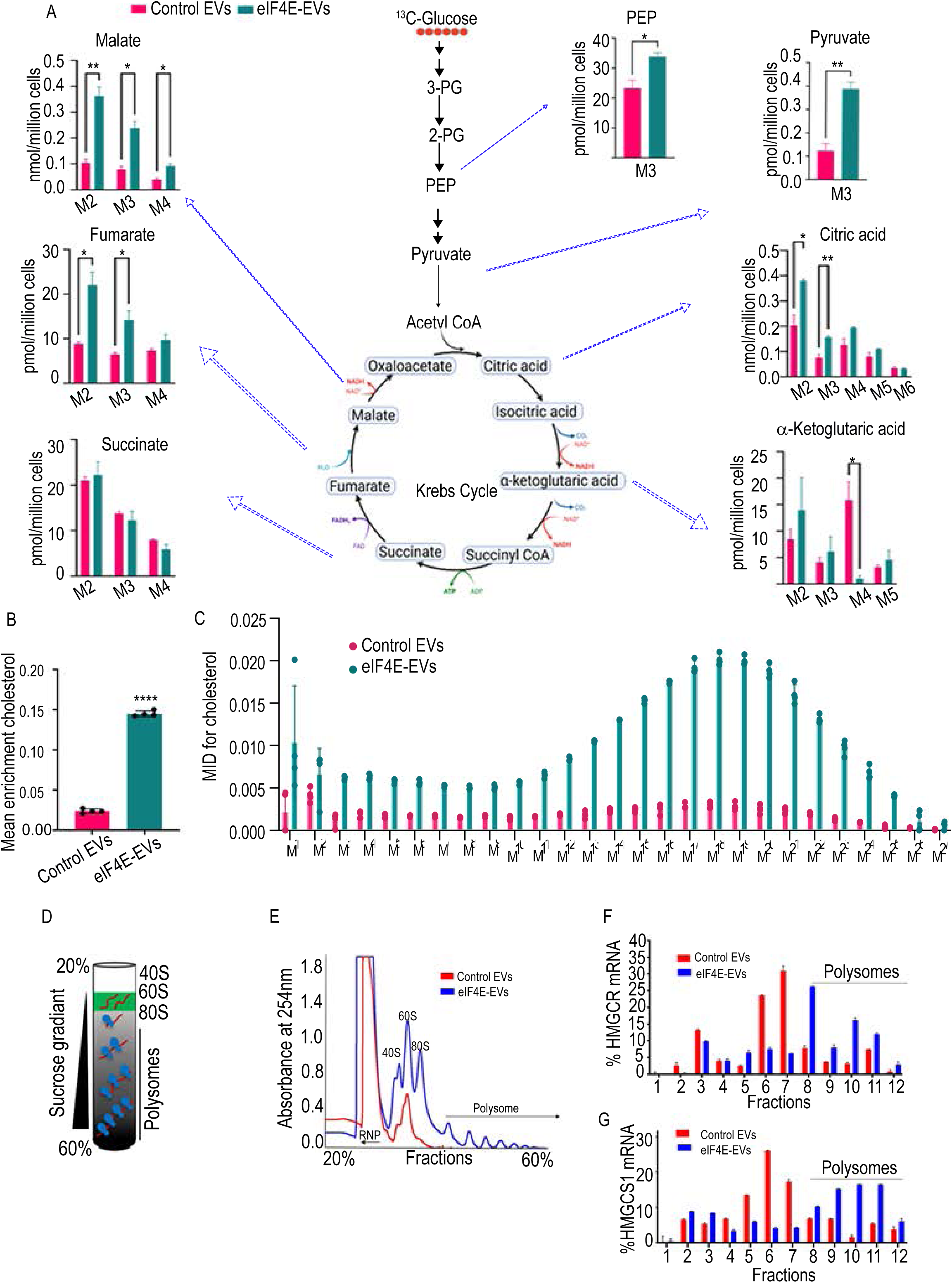
Tumor derived EVs with eIF4E encapsulation enhance de novo cholesterol synthesis in macrophages. (A) Fractional contribution of glucose in central carbon metabolites in control and eIF4E-EVs treated macrophages. n=3 (B) Mean enrichment showing the contribution of uniformly labeled ^13^C glucose in cholesterol. n=4. (C) Mass isotopologue distribution (MID) of cholesterol. n=4 (D) Schema represents polysome profiling. (E) Plot of the absorbance profile of fractions obtained through sucrose gradients to isolate polysomes. (F and G) qPCR shows the enrichment of HMGCR and HMGCS1 mRNA in the isolated fractions bound with free RNPs, monosomes, and polysomes. Data were normalized with the control group. The data are shown as mean ± SEM. *p<0.01, **p<0.05, compared to control by Student’s t-test and one-way ANOVA.

Based on our previous findings, which demonstrated that eIF4E-EVs enhance global protein synthesis (Figure 2O), our objective was to investigate their direct impact on translation efficiency. To achieve this, we treated macrophages with EVs isolated from parental and OVACR8 cells overexpressing eIF4E and prepared cytoplasmic fractions. These fractions were further fractionated using 20%–60% linear sucrose gradient columns (Figure 5D). The absorbance at 254 nm was recorded to profile polysome content, revealing four distinct peaks representing 40S and 60S ribosomal subunits, 80S monosomes, and polysomes from left to right. Notably, our assay detected the presence of elevated polysomal mRNA peaks, indicative of active translation, following the treatment of macrophages with eIF4E-enriched EVs compared to control EVs (Figure 5E). We isolated proteins from gradient fractions, and analyzed via SDS-PAGE and observed a significant increase in the expression level of ribosomal protein RPS6 (S6) in response to eIF4E overexpression mediated by EVs, compared to the control group (Figures S5G-S5H). Subsequently, in another experiment, we determined the abundance of HMGCR and HMGCS1 mRNA within the polysome fractions by qPCR, utilizing mRNA purified from the ribosomal fractions. In line with our previous findings illustrating the translational enhancement induced by eIF4E enriched EVs, we observed an elevated level of HMGCR and HMGCS1 mRNA within the polysome fractions (Figures 5F and 5G).

### 6. Tumor derived EVs enhance tumor growth and elevate PD-L1 expression in macrophages

EVs exhibit the ability to disseminate to various tissues beyond their originating cells. Considering this, our initial investigation focused on the impact of ovarian cancer derived EVs on tumor progression. To accomplish this, C57BL/6 mice were injected with ID8Trp53-^/-^;Brca2^-/-^ cells into the ovary. After 7 days mice were treated with EVs isolated from: (i) ID8Trp53^-/-^;Brca2^-/-^ cells, (ii) ID8Trp53^-/-^;Brca2^-/-^ cells overexpressing eIF4E or (iii) eIF4E knock down ID8Trp53^-/-^;Brca2^-/-^. Mice treated with eIF4E-enriched EVs displayed augmented tumor nodules and ascites volume compared to those injected with control EVs (Figures 6A, 6B, S6A and S6B). Conversely, mice administered with EVs derived from eIF4E KD ID8Trp53^-/-^;Brca2^-/-^ cells exhibited decelerated tumor growth when compared to those receiving EVs from ID8Trp53^-/-^;Brca2^-/-^ cells (Figures 6C,6D, S6C and S6D).

**Figure 6.**
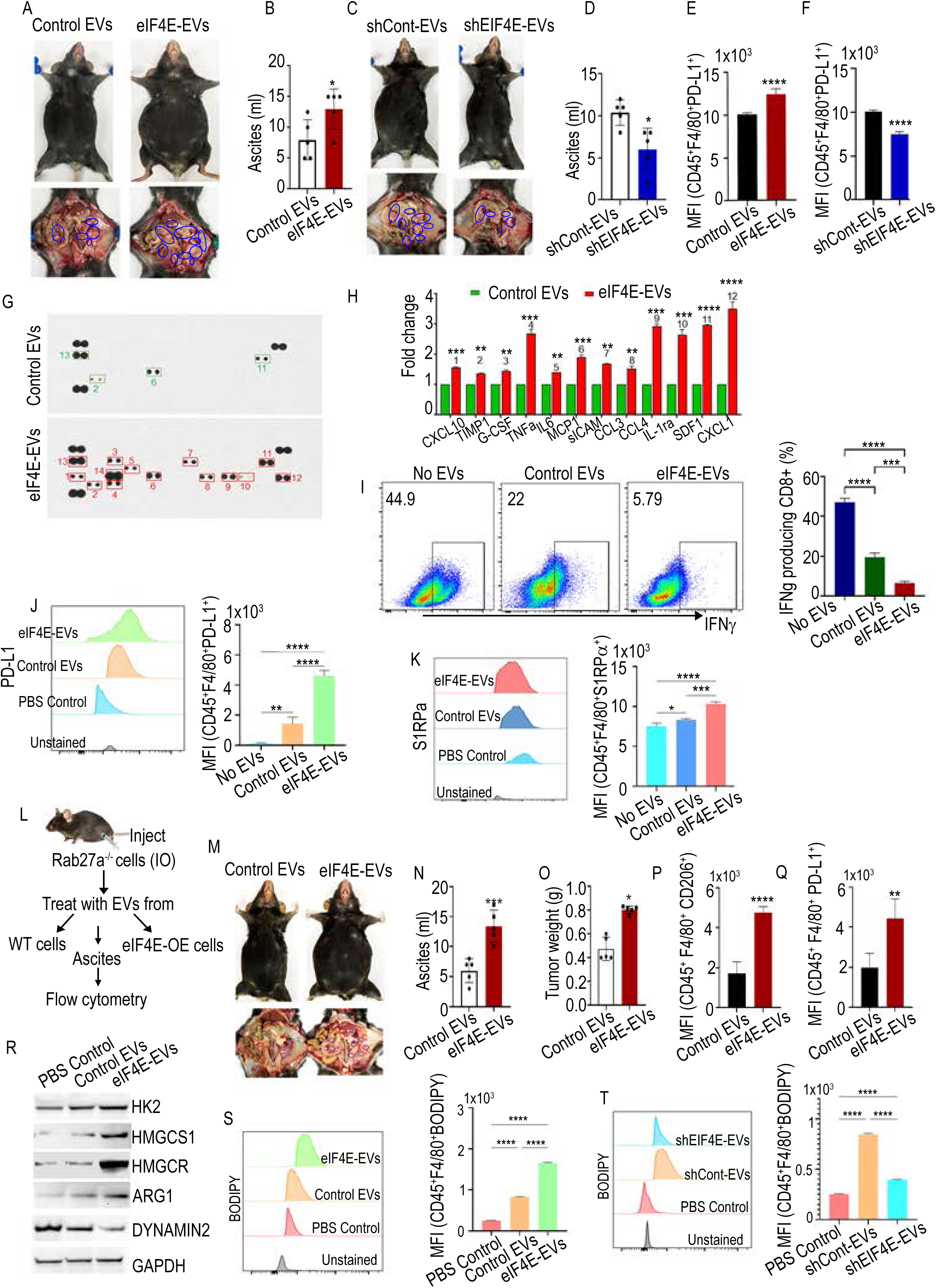
Tumor derived EVs enhance tumor growth and elevate PD-L1 expression in macrophages. (A) Peritoneal cavity of the mouse showing ascites accumulation and tumor locations (blue circles). n=5. (B) Average ascites volume. (C) Peritoneal cavity of the mouse showing ascites accumulation and tumor locations (blue circles). n=5. (D) Average ascites volume. (E and F) PD-L1 expression on macrophages in ascites samples. (G) Representative cytokine array panels of supernatants taken from peritoneal macrophages stimulated with EVs. (H) Representative plots and summarized mean pixel density of each protein are shown. (I) IFN-γ production in activated CD8^+^ T cells. n=3 (J) Expression of PD-L1 on peritoneal macrophages stimulated with EVs. (K) Expression of SIRPα on peritoneal macrophages. (L) Schema for experimental design (M-Q). (M-O) Ascites accumulation and tumor nodules in the peritoneal cavity. (P) CD206 and (Q) PD-L1 on macrophages in ascites. n=3. (R) Western blots showing expression of HK2, HMGCS1, ARG-1 and Dynamin-2 in macrophages. (S and T) Bodipy staining on macrophages isolated from ascites. The data are shown as mean ± SEM. *p<0.01, **p<0.05, ***p<0.001 ****p<0.0001 compared to control by Student’s t-test and one-way ANOVA.

Moreover, our flow cytometry analysis of the ascites demonstrated an enhanced expression of PD-L1 and CD206 with F4/80 in mice treated with eIF4E-EVs (Figures 6E and S6E). While the group treated with eIF4E knockdown EVs exhibited a significant decrease in the expression of PD-L1 and CD206 on F4/80^+^ cells (Figures 6F and S6F). These findings show that exogenously administered eIF4E-packed EVs selectively induce the upregulation of CD206 and PD-L1 expression in macrophages, thereby fostering the emergence of immunosuppressive macrophages within the tumor. To investigate the impact of tumor derived eIF4E-EVs on macrophage functionality, we conducted a cytokine array using the macrophage supernatant. Macrophages stimulated by eIF4E-EVs displayed significant increases in the secretion of TNF-α, IL6, MCP1, CXCL1, IL-1RAa, CCL4, and SDF1 compared to those treated with control EVs (Figures 6G and 6H). To directly assess the impact of eIF4E-EVs-stimulated macrophages on effector T cell function, we pre-treated peritoneal macrophages with either control or eIF4E-EVs before co-culturing them with activated murine T cells. CD8^+^ T cells exhibited a significant reduction in IFN-γ production (Figure 6I), indicating eIF4E-EVs stimulation polarizes macrophages toward an immunosuppressive phenotype.

Consistent with the observed in vivo phenotype, peritoneal macrophages isolated from naïve mice, when subjected to *in vitro* treatment with eIF4E-EVs, exhibited an elevated expression of surface PD-L1 in comparison to control EVs or macrophages alone (Figure 6J)

Apart from the widely recognized PD-1/PD-L1 axis, another pivotal interaction emerging in the field of anti-tumor immunity is the interplay between CD47 and SIRPα. The expression level of SIRPα on eIF4E-EVs stimulated macrophages increased as compared to those treated with control EVs or vehicle (Figure 6K). This data shows that tumor derived EV stimulation skews macrophages towards an immunosuppressive phenotype, involving both direct PD-L1-mediated and indirect cytokine-mediated mechanisms. Subsequently, our objective was to demonstrate that tumor derived EVs are accountable for driving the increased PD-L1 expression in macrophages, thereby facilitating the priming of ovarian cancer metastasis. To address this question, we used CRISPR/Cas9 technology to silence RAB27a, a critical regulatory protein essential for EV secretion ^30, 31^. In brief, we injected RAB27a^-/-^ ID8 cells into the ovary of female C57BL/6 mice resulting in tumors incapable of releasing EVs. Ten days following tumor cell injection, the mice were randomized into three groups and treated with EVs isolated from (i) ID8Trp53^-/-^;Brca2^-/-^ cells, (ii) ID8Trp53^-/-^;Brca2^-/-^ cells that overexpressed eIF4E or (iii) eIF4E knock down ID8Trp53^-/-^;Brca2^-/-^ cells. The mice were closely monitored for tumor progression. We found that the administration of EVs obtained from cells that overexpressed eIF4E increased tumor weight, ascites volume, mice body weight, PD-L1 and CD206 expression on the macrophages compared to EVs derived from parental cells (Figures 6L-6Q and S6G). In another independent experiment, mice injected with EVs derived from eIF4E knockdown cells exhibited reduced tumor burden as well as lower levels of CD206 and PD-L1 expression on macrophages compared to those treated with EVs from ID8Trp53^-/-^;Brca2^-/-^ cells (Figures S6H-S6N).

To investigate the impact of eIF4E-EVs on macrophage metabolism *in vivo*, we isolated TAMs from the ascites through FACS and performed western blotting. In line with our earlier *in vitro* experiments on peritoneal macrophages from naïve mice (Figure 3E), macrophages isolated from eIF4E-EVs treated mice showed increased expression levels of ARG-1, HK2, GLS, HMGCS1, HMGCR, and reduction in Dynamin 2 levels (Figure 6R) compared to macrophages isolated from control EV treated mice. Furthermore, TAMs isolated from mice treated with eIF4E-EVs had the highest lipid content compared to control EVs (Figure 6S). Conversely, TAMs isolated from mice treated with shEIF4E-EVs displayed lower lipid content compared to control EVs (Figure 6T).

Apart from that, there was a significant reduction in the overall tumor burden in the Rab27a^-/-^ group compared to control (Figures S6O-SR). These findings collectively confirm that eIF4E-EVs EVs specifically impact macrophage metabolism to promote tumor growth.

### 7. Inhibition of cholesterol synthesis enhances anti-tumor immune responses

To determine the impact of reducing cholesterol levels in macrophages on their antitumor activity, we injected either control or Hmgcr-KD (shHMGCR)-PMJ2-R cells (Figures 7A and 7B) into Csf1r^-/-^ C57BL/6 mice bearing ID8Trp53^-/-^:Brca2^-/-^ tumors. Mice treated with shHMGCR-PMJ2-R cells displayed reduced ascites volume, tumor weight and mice weight when compared to the control mice (Figures 7C,7D and S7A). Tumor-infiltrating shHMGCR-PMJ2-R cells in tumor-bearing mice exhibited decreased expression of PD-L1 (Figure 7E) and CD206 (Figure 7F), as well as lower cholesterol content when compared to control PMJ2-R tumor-bearing mice (Figure 7G). Given the TAM and TME often exhibit elevated cholesterol levels in comparison to normal cells or tissues, we sought to determine whether reducing cholesterol levels in a tumor or its microenvironment would impact the expression of immune checkpoint markers on macrophages in an in *vivo* setting. To assess the effects of pharmacological inhibition of cholesterol on tumor growth and macrophage activation, we orthotopically injected ID8Trp53^-/-^:Brca2^-/-^ cells into syngeneic C57BL/6 mice and treated with HMGCR inhibitor simvastatin (Figure 7H and S7B).). Simvastatin treatment reduced the ascites and tumor weight when compared to the control mice (Figures 7I and 7J, and S7C). Moreover, simvastatin treatment led to decreased expression of CD206 and PD-L1 on macrophages isolated from ascites, in contrast to control mice (Figures 7K and 7L). Furthermore, CD8+ T cells from simvastatin-treated mice displayed significantly higher Ki67 and the expression level of granzyme B (GZMB), the executor of tumor cell killing, was also increased in CD8+ T cells upon simvastatin treatment (Figure S7D-7F). IHC analysis revealed a marked reduction in Ki67 levels in tumor tissue isolated from mice treated with simvastatin, in comparison to control mice (Figure S7G). These findings highlight the role of cholesterol in immunosuppressive reprogramming and tumor progression. To confirm these results at the cellular level, we assessed the impact of simvastatin treatment on peritoneal macrophages, which exhibited a reduction in PD-L1 expression (Figure S7H). To further gain insights into the underlying mechanism of statin treatment and its impact on PD-L1 downregulation, we employed a transcription factor (TF) array. Macrophages were treated with simvastatin, and nuclear extracts were collected for TF profiling assay. Simvastatin treatment led to a 2.5-fold reduction in the expression of several transcription factors, including EGR, XBP, MZF, NRF-2, OCT-1, Pax-3, MEF1, and Prox1 (Figure S7I).

**Figure 7.**
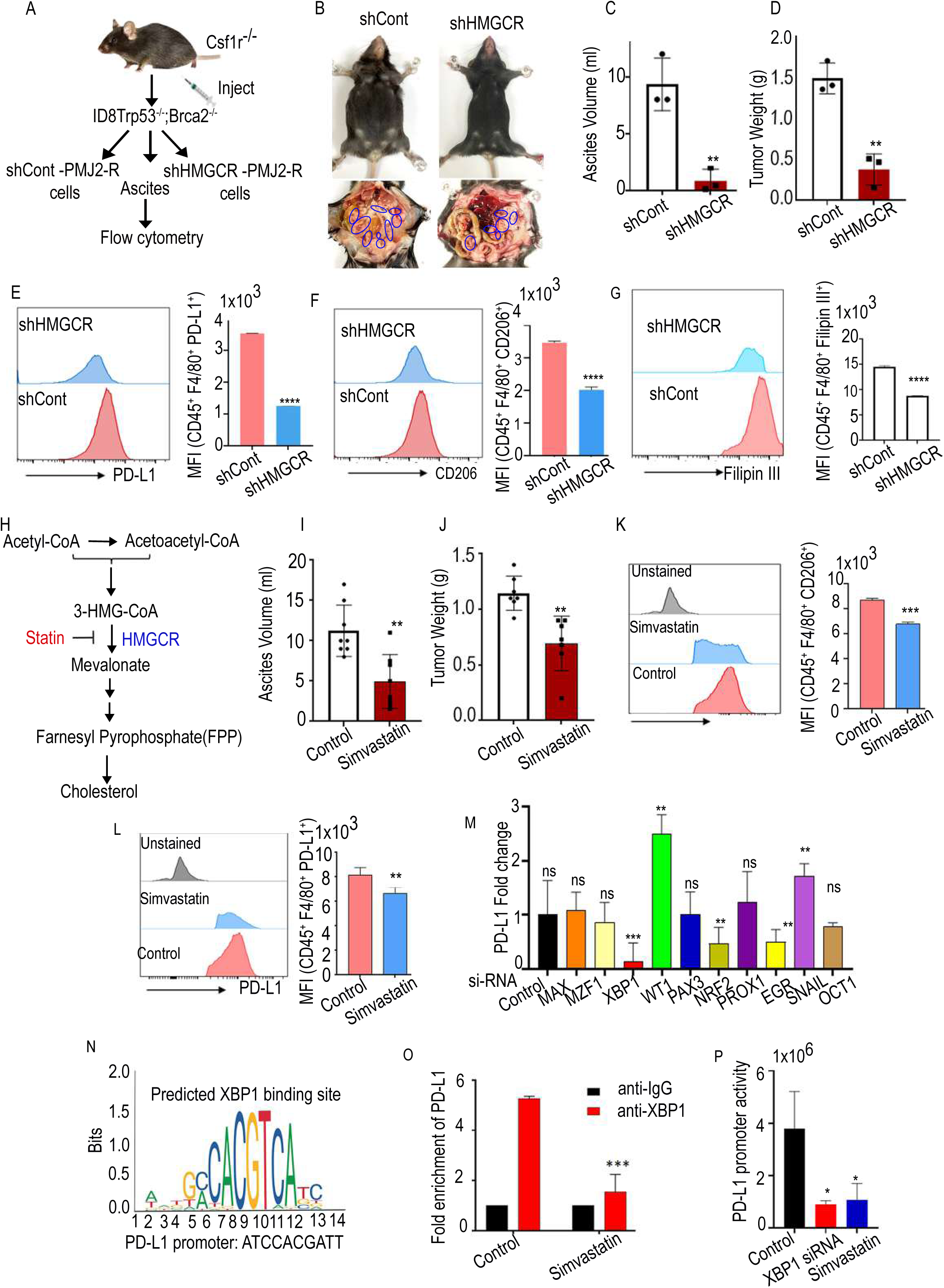
Inhibition of cholesterol synthesis enhances anti-tumor immune responses. (A) Cholesterol biosynthesis pathway. (B) Validation of transcription factor array using qPCR. (C) Predicted XBP1s binding sites on PD-L1 promoters. (D) ChIP analysis of XBP1s binding to PD-L1 promoters in macrophages treated with simvastatin. (E) Dual luciferase analysis of the effect of XBP1 KD and simvastatin treatment on PD-L1 promoter. (F) ascites volume (G) tumor weight. n=7. (H and I) Flow cytometric quantification of the PD L1^+^ and CD206^+^ macrophages in ascites. (J) Schematic workflow. (K) Peritoneal cavity of mice showing ascites accumulation and tumor locations. n=3 (L) ascites volume, (M) tumor weight. (N) PD-L1 and (O) CD206 expression on macrophages in ascites samples. (P) Filipin III staining and flow cytometry of macrophages from ascites samples. Error bars indicate mean ± SEM, *p<0.05, **p<0.01 ****p<0.0001. ns, non-significant.

To further confirm the involvement of these transcription factors, we utilized target-specific siRNAs to knock them down in macrophages, followed by qPCR to assess PD-L1 expression. We found that loss of XBP1 led to a reduction in PD-L1 expression (Figure 7M). Subsequently, we explored the mechanism by which XBP1 regulates the PD-L1 expression on macrophages. Given that XBP1 is a transcription factor ^32^, we analyzed PD-L1 gene promoters and identified potential binding sites for XBP1 (Figure 7N). We then assessed whether simvastatin treatment could attenuate XBP1’s binding to the PD-L1 promoter, by chromatin immunoprecipitation (ChIP) assay. Simvastatin-treated macrophages had reduced binding of XBP1 on the PD-L1 promoter compared with untreated macrophages (Figure 7O). Furthermore, luciferase reporter assay showed that both XBP1 silencing, and simvastatin treatment led to a decrease in PD-L1 promoter activity compared to the control group, indicating that XPB1 activates PD-L1 transcription (Figure 7P).

Collectively, our data establish that cholesterol contributes to the immunosuppressive phenotype of macrophages.

### 8. HMGCR overexpression in TAMs predicts poor prognosis of ovarian cancer patients

Our in vivo experiments demonstrated that elevated HMGCR expression on TAMs plays a pro-tumorigenic role. To assess whether a similar phenomenon occurs in ovarian cancer patients, we obtained tumor tissue from high-grade ovarian cancer patients. Immunofluorescence staining for HMGCR and CD163, a well-established marker of M2-type TAMs, revealed that HMGCR is expressed on CD163+ cells (Figure 8A). Correlation analysis showed that the expression of HMGCR in macrophages was proportional to the number of CD163-positive cells (Figure 8B). We thus speculated that high HMGCR expression was related to intertumoral TAM aggregation in ovarian cancer. To test this hypothesis, we then investigated the potential co-distribution between HMGCR and TAMs in serial sections of ovarian cancer tissue microarrays by IF staining and found increased co-expression of HMGCR and CD163 in malignant high-grade samples (Figure 8C, 8D and S8A),

**Figure 8.**
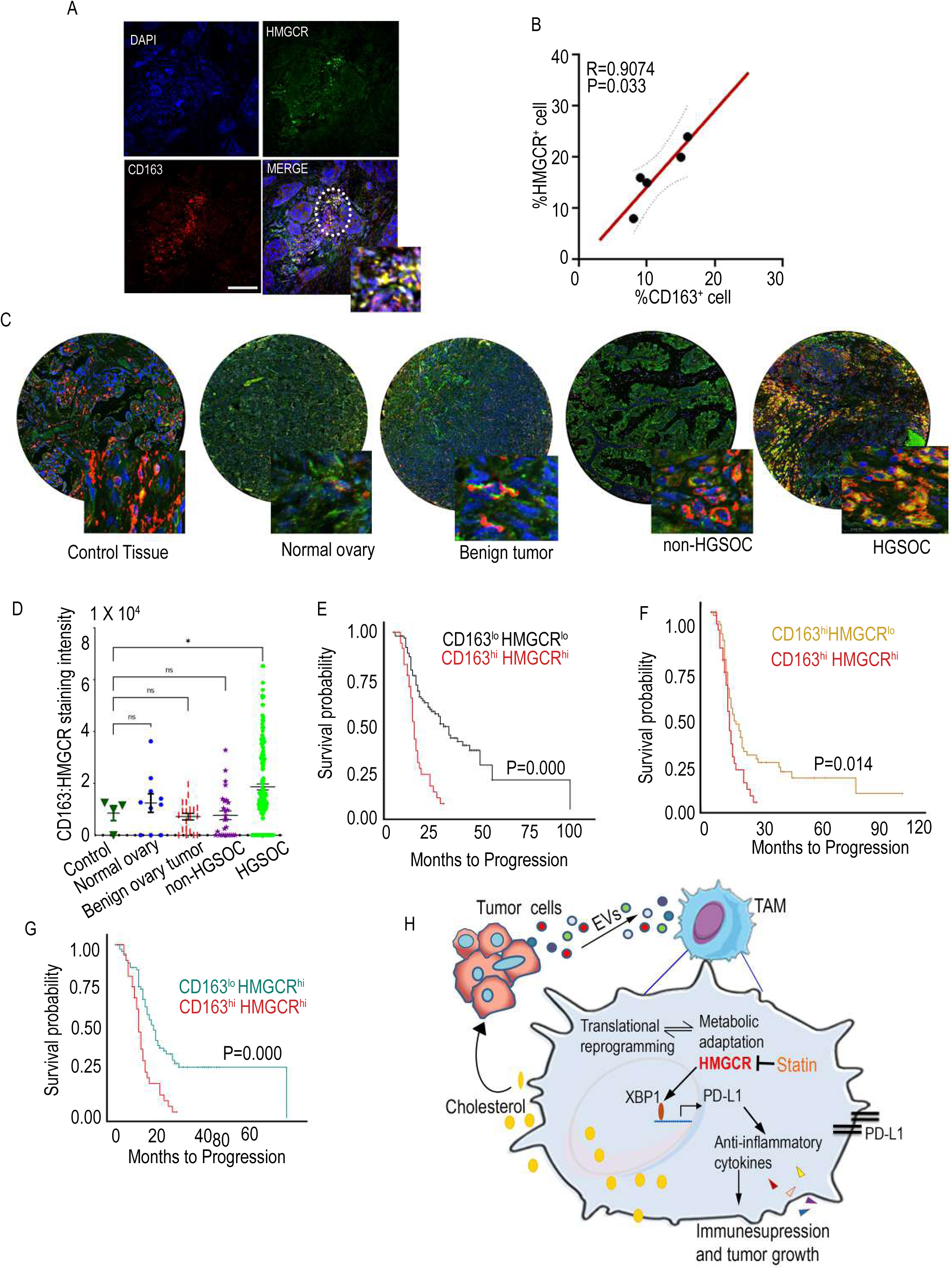
Increased HMGCR expression in TAM predicts poor prognosis. (A) Representative IF staining of HMGCR (Green) and CD163 (red) in human Ovarian cancer tissue. Bar 50 µm. (B) Correlation analysis of CD163^+^/HMGCR^+^ cell proportions in Figure A. (C and D) Co-immunofluorescence staining of HMGCR (Green) and CD163 (red) in human ovarian cancer tissue microarray. (E) Survival curves of high HMGCR and high CD163 expression vs. low HMGCR and low CD163 expression. (F) Survival curves of high HMGCR and high CD163 expression vs. low HMGCR and high CD163 expression (G) Survival curves of high HMGCR and high CD163 expression vs. high HMGCR and low CD163 expression

To further explore the clinical significance of HMGCR upregulation and TAM recruitment in ovarian cancer, survival analyses of associated genes were conducted. Our findings revealed that the expression of CD163 and CD206 alone did not significantly impact overall survival (Figure S8B and S8C). However, patients exhibiting high expression levels of both CD163 and HMGCR had a poorer overall survival compared to those with low expression levels (Figure 8E). Additionally, in patients with high CD163 expression better survival outcomes were observed in those with low HMGCR expression (Figure 8F). Similarly, in patients with high HMGCR expression better survival outcomes were observed in those with low CD163 expression (Figure 8G). Survival analyses of CD206 and HMGCR expression also yielded consistent results (Figure S8E-S8G). Collectively, these findings suggest that HMGCR-positive TAMs are linked to worse prognosis in ovarian cancer patients. This supports the notion that HMGCR signaling is elevated in TAMs, potentially contributing to the initiation and progression of ovarian cancer.

## Discussion

Extracellular vesicles produced by cancer cells serve as a distinctive mode of intercellular communication, exerting influence on cell growth and survival, and helping shape the tumor microenvironment. These EVs carry various cargo, including mRNA, miRNA, proteins, and transcription factors, which can modulate gene expression and cellular functions in both normal and cancer cells. Studies have established a direct correlation between the presence of EVs and gene regulation ^33, 34^. While the contents of EVs are characterized as both diagnostic and prognostic biomarkers, the impact of EVs on tumor resident immune and stromal cells are not well characterized. Importantly, our study provides evidence that how EVs dysregulate function of T cells and create an immunosuppressive environment due to translational adaptations in tumor resident macrophages.

For further characterizing the impact of the contents in EVs, we have used an unbiased mass spectrometric approach to characterize the proteins expressed in EVs. Strikingly, we found that the tumor derived EVs exhibit high levels of translation initiation factor eIF4E. Mechanistically, we demonstrated that eIF4E interacts with UEV domain in TSG101 for encapsulated into EVs.

eIF4E protein is a critical part of the eukaryotic translation initiation factor 4F complex, which recognizes the 7-methylguanosine cap structure at the 5’ end of messenger RNAs and recruiting ribosomes to the 5’-cap structure. eIF4E, a cap-binding protein, collaborates with proteins like eIF4A and eIF4G to bind mRNA, facilitating ribosome recruitment and translation initiation ^35^. In brief, the association of eIF4E with the 4F complex is considered the rate-limiting step in translation initiation ^36,37^. Notably, increased expression of eIF4E has been linked to various oncogenic activities in ovarian cancer cells, including proliferation, migration, invasion, and chemosensitivity ^37, 38^. Supporting the notion that eIF4E is associated with oncogenic functions, our data demonstrates how eIF4E influence translation process in macrophage through the uptake of EVs loaded with eIF4E.

Metabolic interplay among immune cells, stromal cells, and cancer cells significantly impacts tumor evasion from immune surveillance, fostering cancer progression and metastasis ^39, 40^. TAMs, which are associated with all stages of neoplastic progression in patients, comprise a large proportion of the immune infiltrate in malignancies. Ovarian cancer cells are known to promote cholesterol efflux in TAM by inducing IL4 mediated arginase programming ^41^.

We elucidate how eIF4E-packaged EVs affect metabolic profiles in macrophages within the tumor microenvironment. Importantly, eIF4E uptake in macrophages induced the translation of ribosomal proteins and key enzymes HMGCR and HMFCS1 involved in cholesterol synthesis. Metabolic analysis also revealed that eIF4E uptake in macrophages enhanced nucleotide biosynthesis, pentose phosphate pathway activity, increased the levels of enzymes involved in glucose, glutamine, and cholesterol metabolism and elevated levels of amino acids essential for protein synthesis.

We further observed that the metabolic adaptations promote the immunosuppressive characteristics of macrophages such as high expressions of arginase and PD-L1. Importantly, our study demonstrates that inhibiting EV secretion reduces PD-L1 expression on TAMs in ovarian tumor-bearing mice, highlighting the role of eIF4E in modulating PD-L1 expression and influencing metastasis in ovarian cancer.

Cholesterol-lowering drugs, particularly statins, have broad clinical implications ^42-44^. Our study shows the potential therapeutic implications, as cholesterol-lowering drugs, particularly statins, exhibit anti-cancer properties by reducing PD-L1 expression in cancer cells and modulating cholesterol metabolism in macrophages via the XBP1 transcription factor. Overall, our study uncovers an overlooked oncogenic mechanism involving eIF4E-packaged EVs that upregulate cholesterol synthesis in macrophages, fostering immune suppression within the tumor microenvironment.

To determine the if HMGCR expression in TAM are associated with poor prognosis and tumor progression, we checked the expression of HMGCR and CD163 in ovarian cancer patients. We found a positive correlation between HMGCR and M2 TAM-related molecules in ovarian cancer. The findings emphasize the significance of targeting cholesterol metabolism in macrophages to enhance the efficacy of cancer immunotherapies.

## RESOURCE AVAILABILITY

### Lead contact

Further information and requests for resources and reagents should be directed to and will be fulfilled by the lead contact, Sunila Pradeep.

### Materials availability

This study did not generate new unique reagents.

### Data and code availability

This paper does not report the original code. Any additional information required to reanalyze the data reported in this paper is available from the lead contact upon request.

## EXPERIMENTAL MODEL AND SUBJECT DETAILS

### Patient Samples

After informed consent, peripheral blood was collected from ovarian cancer patients and healthy individuals according to an Institutional Review Board approved protocol at Medical College of Wisconsin. All the clinical samples we used are from adult females.

### Cell lines

Human ovarian cancer cells, HeyA8 was purchased from the Characterized Cell Line repository at MD Anderson Cancer Center, Houston, TX, USA. IOSE-21, OVCAR4, A2780, OVCAR5, and OVCAR8, were purchased from the National Cancer Institute (NCI). NIH-OVCAR3 was purchased from ATCC/PBCF repository. THP1, PMJ2-R and RAW264.7 cell lines were purchased from ATCC. OVSAHO and Kuramochi cells were received from Taru Muranen at Beth Israel Deaconess Medical Center, Boston, Massachusetts, USA. BR-Luc murine cell lines were received as a kind gift from Sandra Orsulic (University of California, Los Angeles, CA).

ID8Trp53^-/-^;Brca2^-/-^ murine ovarian cancer cell line was a kind gift from Dr. Iain A. McNeish, Wolfson Wohl Cancer Research Centre, Institute of Cancer Sciences, University of Glasgow, Glasgow, United Kingdom. HEK293-T cell line was procured from Thermo Fisher Scientific. LP3 and Untransformed fibroblast cell line (GM07213) was procured from Coriell Institute for Medical Research, NJ, USA. RF24 was received from Dr. Arjan W. Griffioen, VU University Medical Center, Amsterdam, Netherlands. Ovarian surface epithelial cells (OSE) were cultured by scraping the surface epithelium of normal ovarian tissues obtained from two patients with benign gynecologic pathology. ID8Trp53^-/-^;Brca2^-/-^ cells were cultured were cultured in Dulbecco’s modified Eagle’s medium (DMEM) (Sigma Aldrich) with 4% FBS (Atlanta Biologicals, GA, USA), and 1x ITS. NIH OVCAR3 was cultured in RPMI-1640 Medium with 10% FBS (Atlanta Biologicals, GA, USA), and HEPES (ATCC). OSE cells were cultured in MEGM media (Lonza, Basel, Switzerland) supplemented with 15% FBS. Fibroblast cells were cultured in Eagle’s minimum essential medium with Earle’s salts and non-essential amino acids and 10% non-heat inactivated FBS. LP3 cells were cultured in medium 199 and MCDB (1:1) supplemented with 15 % non-heat inactivated FBS, 10ng/ml EGF and 0.4µg/ml hydrocortisone. THP1, and Raw264.7 cells were cultured in RPMI-1640 medium with 10% FBS and HEPES. RF24 cells were cultured in Eagle’s minimum essential medium supplemented with 10% FBS, non-essential amino acids and modified Eagle medium vitamins. All other cell lines were cultured in Dulbecco’s modified Eagle’s medium (DMEM) (Sigma Aldrich) supplemented with 10% FBS and 1% Pen-strep (Thermo Fisher Scientific Inc., Waltham, MA, USA) at 37°C in a humidified incubator with 5% CO_2_. The cell lines were authenticated by short tandem repeat (STR) profiling (IDEXX BioAnalytics) and tested for Mycoplasma using MycoSensor PCR Assay kit (Agilent, Santa Clara, CA)

### Mice

C57BL/6 black female mice were purchased from Envigo, B6.Cg-Csf1r<tm1.2Jwp>/J female mice were purchased from the Jackson Laboratory. All experiments complied with protocols approved by the Institutional Animal Care and Use Committee at the Medical College of Wisconsin.

## METHOD DETAILS

### EV isolation, characterization, and quantification

Supernatant was collected from ovarian cancer cells and non-cancerous epithelial cells grown to 80% confluency in serum free media for 48-72h. EVs were purified and characterized from supernatant as previously described^2, 45^. To purify EVs using sucrose gradient, resulting supernatant underwent ultracentrifugation at 4°C to pellet EVs. These EVs were resuspended in PBS and layered onto a sucrose gradient, followed by centrifugation. The isolated EV pellets were lysed, and total EV protein concentrations were determined by BCA Protein Assay kit (Thermo Fisher Scientific, Santa Clara, CA). The EV size distribution and concentrations were determined by Nanoparticle Tracking Analysis (NTA), using the NanoSight LM10 instrument (Malvern Panalytical, Malvern, UK) with a 488nm laser and NTA3.1 software. Purified EVs were fixed in 2% paraformaldehyde for transmission electron microscopy (EM) following the protocol of Bulreigh et. Al. ^46^ and examined in a Hitachi H600 TEM (EM facility at the Medical College of Wisconsin USA). To assess the uptake of EVs by different cells EVs were labeled using DIL red fluorescent labeling dye (Invitrogen). The DIL-labeled EVs were then incubated with cells at 37 °C. The uptake of labeled EVs was assessed using confocal laser scanning microscope (LSM 510; Zeiss, Oberkochen, Germany).

### Protein extraction and trypsin digestion

EV pellets were resuspended in a solution containing 40% Invitrosol, 20% acetonitrile, and 100 mM ammonium bicarbonate followed by sonication. Total protein concentrations were measured, and cysteines were reduced and alkylated. Proteins were then digested with LysC and trypsin enzymes. Peptides were cleaned up using paramagnetic particles, and their concentrations were determined. Samples were diluted using acetonitrile and formic acid, with the addition of a peptide retention time calibration mixture.

### Mass spectrometry analysis

Peptides were dissolved and separated using a NanoElute ultra-high-performance liquid phase system. Each sample underwent analysis on a Thermo Scientific Orbitrap Fusion Lumos MS via 3 technical replicate injections using data-dependent acquisition (DDA). MS data were analyzed using the Proteome Discoverer 2.4 platform, referencing the Swiss-Prot_Human with isoforms version from 2019-05-01. Differentially expressed genes were assessed within biological pathways using the Ingenuity Pathway Analysis software (IPA; Ingenuity Systems Inc). Fisher’s exact test calculated the p value to determine a potential significant association between differentially expressed proteins and specific functional categories. A p value < 0.05 was deemed statistically significant. Identified proteins were cross-referenced with exosome data available from the ExoCarta database (http://www.exocarta.org).

### Immunohistochemistry and Immunofluorescence

Slides were dewaxed in xylene and rehydrated through graded ethanol to distilled water. Antigen retrieval for the slide specimens were performed using IHC-Tek epitope retrieval solution and steamer set (IHC World, LLC.). The slides were then immersed in 3% H_2_O_2_ for 10min to quench endogenous peroxidase followed by blocking with 10% goat serum for 1h. Slides were incubated with primary antibodies, and then secondary antibodies. Slides were stained with DAB and hematoxylin, dehydrated, mounted, then cover-slipped. For immunofluorescence, the sections were incubated for 1 hour in the dark with fluorochrome-conjugated secondary antibodies. Nuclei were stained using Antifade Fluorescence Mounting Medium with DAPI. Slides were digitally scanned using Panoramic 250 FLASH III scanner (3D HISTECH ltd. Version 2.0) and, the slide Viewer software (3D HISTECH ltd. Version 2.0) was used to view and analyze images.

### Western blotting

An equal amount of total protein was resolved on precast 4%–12% SDS-PAGE gels (Biorad, Hercules, CA, USA), and the protein was transferred onto PVDF membranes as previously described ^2^. Membranes were incubated with the desired primary and secondary antibodies. Protein expression was detected with a chemiluminescence kit.

### siRNA transfection

Predesigned siRNAs were purchased from Dharmacon, Lafayette, CO.Negative siRNAs universal controls (siCont) were obtained from Sigma-Aldrich, which also supplied us with predesigned siRNAs. Transfections were performed using the Lipofectamine RNAiMAX transfection reagent (Thermo Fisher Scientific Inc., Waltham, MA). At 48h post transfection, cells were harvested for further analysis.

### Co-immunoprecipitation GFP Trap

OVCAR8 cells stably expressing either GFP fusion proteins or TSG101 variants were washed with PBS and lysed by adding lysis buffer supplemented with protease inhibitor. Cleared lysates were incubated with GFP-nanobody agarose (GFP-Trap, Chromotek). The beads were washed with ice-cold Co-IP washing buffer and proteins were eluted in sample buffer by boiling at 95°C for 10min followed by western blotting.

### Proximity ligation assay

The DuoLink® In Situ Red Starter Kit Mouse/Rabbit (Sigma-Aldrich) was used to detect proximity between eIF4E/eIF4A1and TSG101 proteins according to the manufacturer’s protocol. Fluorescence images were acquired with a confocal laser scanning microscope.

### SUnSET assay

Macrophage (THP1-derived), endothelial (RF24), mesothelial (Met-5A, LP3) and fibroblast (GM07213) cells were incubated with EVs isolated ovarian cancer cells. After 24h, cells were incubated with puromycin for different time points. Cells were lysed using 1x RIPA buffer lysates and run on an SDS-PAGE gel, transferred to a nitrocellulose membrane for western blotting with anti-puromycin antibody Puromycin staining in lanes was measured using chemiluminescence kit.

For flow cytometry, THP-1-derived macrophages were incubated with EVs and treated with CHX for 1h followed by incubation with puromycin. Cells were washed with PBS to remove residual puromycin followed by staining of cells using Alexa fluor-488 anti-puromycin antibody.

### Polysome fractionation by sucrose gradients

Polysome fractionation was performed following a previously published protocol ^47, 48^. In brief, THP1-derived macrophages were incubated with EVs, and total lysate was prepared. Approximately 10-15ml of lysate were layered over 20% – 60% cold sucrose gradients. Gradients were centrifuged in a Beckman SW28 rotor for 2h at 4°C. After centrifugation, 12 equal-sized fractions were collected using polysome fractionator. For western blotting, fractions were precipitated and mixed with SDS sample buffer. For qPCR, total RNA was isolated from the fractions by mixing with phenol:chloroform:isoamyl alcohol (Sigma-Aldrich) and then RNA was analyzed by qRT-PCR.

### *In vivo* Extracellular vesicle treatment

Rab27a^−/−^ ID8 cells were injected orthotopically into the ovary of C57BL/6 mice. Seven days post-injection, EVs isolated by ultracentrifugation from culture media of ID8Trp53^-/-^;Brca2^-/-^ cells, ID8Trp53^-/-^;Brca2^-/-^ eIF4E KD and ID8Trp53^-/-^;Brca2^-/-^ cells with eIF4E overexpressed were injected into mice via the tail vein thrice a week until the control mice became moribund.

### *In vivo* macrophage treatment

For adoptive transfer, Csf1r^-/-^ mice (3-5/group) were injected intraperitoneally with ID8Trp53^-/-^;Brca2^-/-^ cells. After 10 days, tumor-bearing mice were treated with immortalized peritoneal macrophages (PMJ2-R) twice a week. After 7 weeks, mice were sacrificed, tumor weight and ascites volume were recorded.

### Isolation and culturing of peritoneal macrophages

Peritoneal macrophages were isolated from black mice by peritoneal lavage as described previously ^49^. Briefly, cold PBS containing 1% FBS was injected into the peritoneal cavity and extracted after gentle agitation. The peritoneal cell suspension was centrifuged, and cells were resuspended in complete RPMI-1640 supplemented with 10% FBS (Atlanta Biologicals), and 1% Penicillin-Streptomycin Solution (Corning). Cells were incubated at 37°C for 2h to allow macrophages to adhere to the plate. Floating cells were removed by two subsequent washes with PBS. For cell sorting, peritoneal macrophages were stained with CD45 and F4/80 antibody (Biolegend) at 4°C for 1h in the dark. Samples were then washed and resuspended in FACS running buffer.

### Quantitative real time-PCR (qRT-PCR)

Total RNA was isolated from the cells using the RNeasy Mini Kit (Qiagen, Valencia, CA, USA), and first-strand cDNA was transcribed using iScript reverse transcription supermix (Biorad, Hercules, CA, USA). qRT-PCR was performed using CFX Connect Real-Time PCR systems (Biorad, Hercules, CA, USA) and SYBR Premix Ex Taq II (Biorad, Hercules, CA, USA).

### Stable cell line construction

eIF4E/eIF4A1 overexpressed stable cell lines were developed using the lentivirus vector from Genecopoeia following the manufacturer’s instructions. The stable cell lines were selected by adding puromycin into the culture medium. Lentiviral knockdown cell lines were generated as previously described ^50^. Briefly, 293T cells were co-transfected with these plasmids: psPAX2, pMD2.G and pLKO.1 containing shRNA sequence targeting eIF4A1, eIF4E and HMGCR.

sh-eIF4E – 5’ CGATTGATCTCTAAGTTTGAT3’; 5’CCGAAGATAGTGATTGGTTAT3’

sh-eIF4A1-5’GATCTATGACATATTCCAGAA3’; 5’CGAGAGTAACTGGAACGAGAT3’

sh-HMGCR-5’ CCCACAAATGAAGACTTATAT; 5’GAACTTTGCAATCTAAGTTTA

Two days following co-transfection, competent lentivirus particles were collected. ID8Trp53^-/-^; Brca2^-/-^ cells were infected with sh-eIF4E and sh-eIF4A1 lentivirus for 48h. PMJ2-R cells were infected with sh-HMGCR lentivirus. Stable knockdown cell lines were selected with puromycin in the cell culture medium.

### CRISPR/CAS9 knockdown of genes

RAb27a genes knockout ID8 cells were engineered by Synthego (CA, USA) using CRISPR/CAS9 technology. The following guide sequence were used:

RAB27a-CCUGCAGUUAUGGGACACGG

### Transcription factor array

THP1-derived macrophages were treated with 10µm simvastatin for 24h and nuclear protein was extracted from treated and untreated cells. We measured the activity of 96 transcription factors in protein extract using the TF Activation Profiling Plate Array II (Signosis) according to manufacturer’s instructions, Relevant TFs were selected by the fold change (>2 fold) between simvastatin treated and untreated THP1-derived macrophages.

### Luciferase reporter assay

Plasmid vector containing the promoter of PD-L1 was purchased from GeneCopoeia and transfected into cells by Lipofectamine-2000 reagent. The cells were then transfected with XBP1 siRNA or treated with simvastatin for 24h. Culture medium was then collected and centrifuged at 2000×g for 10min at 4°C, and 10µl samples of supernatant were transferred into white-walled 96-well plates in triplicate. Luciferase intensity in each well was immediately measured, using a luminometer, as described in the Secrete-Pair Gaussia LuciferaseAssay Kit.

### ChIP assay

We used the CiiDER tool to determine if the PD-L1 promoter contains a binding site for the transcription factor XBP1. THP1-derived macrophages were treated with 10uM simvastatin for 24h. We performed the ChIP assay using SimpleChIP® Enzymatic Chromatin IP Kit (Cell Signaling Technology by adding XBP1 antibody to both samples. Real-time PCR were used to analyze the amplification of region when XBP1 binds to the PD-L1 promoter Quantitative ChIP-PCR primers:

PD-L1-F:5’-ACACACACACACACACCTAC-3’

PD-L1-R:5’CCCCTTCAAGGTGACTGAACAT-3’

### Cytokine measurement

Peritoneal macrophages from C57BL/6 naïve mice were treated with EVs derived from the supernatants of the ID8Trp53^-/-^;Brca2^-/-^, and eIF4E overexpressed ID8Trp53^-/-^;Brca2^-/-^. After incubation, supernatant was isolated from cultured macrophages, centrifuged at 6000rpm for 5 min to remove cellular debris and transferred to a new Eppendorf tube. The Cytokine array was performed using the Proteome Profile Mouse Cytokine Array Panel A (R&D Systems, Minneapolis, MN) according to the manufacturer’s instructions.

### Mouse T cell isolation

Spleen was isolated from C57BL/6 mice and CD8 T cells were isolated using MojoSort™ Mouse CD8 T Cell Isolation Kit (Biolegend). The cells were stimulated with Dynabeads™ Mouse T-Activator CD3/CD8 kit (Gibco). T cells were co-cultured with macrophages pre-treated with EVs at a 15:1 ratio. Macrophages were washed prior to T cell addition to remove residual EVs and recounted. Following culture, cells were surfaced stained with CD8/BV605 for 30min. Cells were then fixed (Fixation Buffer, Biolegend) and permeabilized (Intracellular Staining Perm Wash Buffer, Biolgend) prior to overnight intracellular staining with IFN-γ.

### Flow cytometry

The single-cell suspension was first incubated with FC block, then live-dead staining was performed, and the cells were washed with stain buffer. Cell surface antigens were stained by co-incubation with antibodies for 30min on ice followed by washing with stain buffer. Staining of intracellular antigen was performed with a BD Pharmingen kit according to the manufacturer’s protocol. Flow cytometry was performed on a Fortessa X20 and Cytek Aurora, and data were analyzed by FlowJo software.

### Stable Isotope Labeling by/with Amino acids in Cell culture (SILAC) based mass spectrometry

SILAC RPMI (Pierce Biotechnology) was supplemented with 10% dialyzed fetal bovine serum (Thermo Scientific, Waltham, MA, USA), 1% streptomycin/penicillin. The heavy medium was supplemented with ^13^C_6_ L-arginine and ^13^C_6_ ^15^N_2_-L-lysine. The light medium was supplemented with normal L-arginine and L-lysine. For SILAC experiments, THP1-derived macrophages were grown in parallel in either light or heavy media for 5 days, with media replacement every 24h. Cells grown in light medium were incubated with control EVs and cells with heavy medium were incubated with eIF4E-EVs for 24h.

The SILAC-labeled cell pellets were lysed and analyzed on a Thermo Scientific Orbitrap Fusion Lumos MS via 3 technical replicate injections using a data-dependent acquisition (DDA) as described earlier.

Peptides and proteins were both filtered for 0.01 FDR, proteins a minimum two unique peptides, precursor abundance based on area, unique and razor peptides used for quantification, normalized on total peptide amount. Quantified proteins with ≥ 2 and ≤ 0.5-fold change were selected and clustered by biological functions, pathway and network analysis using Ingenuity Pathway Analysis (IPA) software (www.ingenuity.com) for bioinformatics analysis. A heat map for differentially expressed proteins was generated using heatmapper (http://www.heatmapper.ca/).

### Metabolism assays

#### Seahorse XF cell Mito stress test

An XF96e Analyzer (Agilent Technologies, Santa Clara, CA) was used to measure bioenergetic function in isolated peritoneal macrophages and THP1-derived macrophages as described earlier ^51^. In brief, cells were stimulated with control and eIF4E-EVs overnight. For all bioenergetic measurements, the culture media was changed 1h prior to the assay run to unbuffered Agilent Seahorse XF RPMI Medium supplemented with D-Glucose Pyruvate and glutamine Three basal OCR measurements were recorded prior to injection of oligomycin. After recording the oligomycin-sensitive OCR, FCCP-sensitive rates were recorded. Finally, antimycin A/rotenone was injected to inhibit electron flow through the electron transport system.

### Seahorse glycolytic stress assay

Cells were stimulated with control and eIF4EEVs overnight. The culture media was changed 1h prior to the assay run to unbuffered Agilent Seahorse XF RPMI Medium supplemented with glutamine Three basal ECAR measurements were recorded followed by an injection of a saturating level of glucose, measuring a glucose-induced response. This was followed by Oligomycin A, an ATP synthase inhibitor, which inhibits mitochondrial ATP production and shifts the energy production to glycolysis, revealing the cellular maximum glycolytic capacity. Finally, 2-deoxyglucose, a glucose analog, was injected to inhibit glucose binding to hexokinase, thus confirming extracellular rates to be a product of glycolysis.

### *In vitro* metabolism assays

Peritoneal macrophages and THP1-derived macrophages were stimulated with control and eIF4E-EVs overnight. Cells were then washed and incubated in glucose free RPMI for 30min before 2-NBDG, 100-200μg/ml in glucose-free medium (Caymen-600470) was added. Cells were incubated with 2-NBDG for 2h before washing. 2-NBDG taken up by cells was detected with fluorescence plate reader at excitation and emission wavelengths of 485nm and 535nm, respectively.

### Cholesterol Content Measurement

Peritoneal macrophages and THP1-derived macrophages were stimulated with control and eIF4E-EVs overnight. After incubation lipid was extracted and total cellular cholesterol content was measured in macrophages using Amplex Red Cholesterol Assay kit (Invitrogen) according to the manufacturer’s instructions. EVs treated macrophages were also co-cultured with tumor cells. After incubation tumor cells were washed and cholesterol content was measured in tumor cells. For the detection assay, cells were stained with Filipin III and then analyzed by flow cytometry.

### Metabolite extraction

Steady state isotope tracing analysis using mass spectrometry was performed as reported earlier ^52-54^. Briefly, THP1-derived macrophages were seeded in 6-well plates overnight and replaced with medium containing U-^13^C_6_ glucose or U-^13^C_5_ glutamine. Cells were treated with control EVs and eIF4E-EVs and metabolites were extracted.

#### LC-MS/MS for untargeted metabolomics

Dried extracts were reconstituted in methanol/water (50:50), sonicated and filtered. 5µL for the sample was injected for untargeted analysis on Kinetex F5 column (2.1 x 150mm, 2.6µm) in both positive and negative ion modes. The column was kept at 30°C and the flow rate was 0.2ml/min. The ScieXOS software was used for peak integration and analysis.

#### GC-MS for cholesterol

Dried cholesterol samples were derivatized with TMS and incubated at 37°C for 30 min. Samples were analyzed using Agilent 7890 GC equipped with a 30-m HP-5MSUI capillary 1246 column connected to an Agilent 5977B MS in scan mode. 1-2µL of sample was injected at 270°C with helium as the carrier gas at 1ml/min flow. The temperature gradient was maintained at 260°C for 3 min, raised to 280°C for 15min, increased to 325°C for 15min and held for 15 min. The quadrupole was operated at 150°C.

#### GC-MS for polar metabolites

Dried samples were derivatized using methoxyamine hydrochloride (MOX, FisherScientific, PI45950) followed by incubation at 45°C for 30 min with constant shaking. Next, 30μL of MBTSTFA+1% TBDMCS (Sigma Aldrich, M-108-12435X1ML) was added, and the samples were incubated at 45°C for 1 h. 1-2µL of the derivatized sample was injected at 270°C with helium as the carrier gas at 1ml/min flow. Temperature gradient method was set up as 100°C for 1 min, raised at 3.5 °C/min to 255°C, increased to 320°C at 15 °C/min and held for 3min (method total time 52.6min). MS detector was operated in scan mode (70–600 m/z). A calibration curve was run for polar metabolite standards (0-10nmoles).

### Quantification and Statistical Analysis

Statistical analysis was done using GraphPad Prism 9.5.0 and presented as mean values ± SEM. For comparison of two groups, Student’s t-test was performed to calculate p values. For more than two groups, one-way ANNOVA and post-hoc Dunnett’s multiple comparison tests were used to compare data from control and each test group. Throughout all figures: ∗p < 0.05, ∗∗p < 0.01, ∗∗∗p < 0.001, and ∗∗∗∗p < 0.0001. Significance was defined at p < 0.05.

## Acknowledgments

This work was supported in part to S.P. by, the Women’s Health Research Program (WHRP), the Department of Defense (DoD W81XWH 21 1 0361 and W81XWH-21-1-0138), and NCI R01CA258433. P.C.□R. was supported by DoD W81XWH 21 1 0365, NCI R01CA229907, and Linda G. and Herbert J. Buchsbaum Endowment. S.M was supported by ovarian cancer research alliance (OCRA, Grant ID#: MIG-2023-2-1016), We thank Research Core services in Medical College Wisconsin Cancer Center, Children’s Research Institute, Versiti BRI and tissue bank from Froedtert hospital Milwaukee.

## Authors contribution

S.M., P.C.□R., and S.P. initiated the study and designed the experiments. S.M. performed most of the experiments, statistical analysis, prepared figures and drafted the manuscript. M.N, O.A, N.M, A.S helped with LC-MS and GC-MS experiments and analysis. S. W.T. performed all the bioinformatics and computational analysis for this study. I.P.K, S.K. J.G, P.G, M.S, A.G and C.D. assisted on animal experiments and in vitro experiments. P.C.R. provided scientific feedback and assisted with translational experiments. D.N provided scientific feedback and assisted in designing and executing LC-MS and GC-MS experiments. S.P established collaborations and allocated funding for the work. SM prepared the manuscript with S.P., All authors have read and approved the article.

