## Supplementry data for "Eukaryotic Translation Initiation Factor Loaded Extracellular Vesicles Promotes Macrophage Cholesterol Metabolism in ovarian cancer"

**Supplementary Figures**

**Sup Fig-1.**

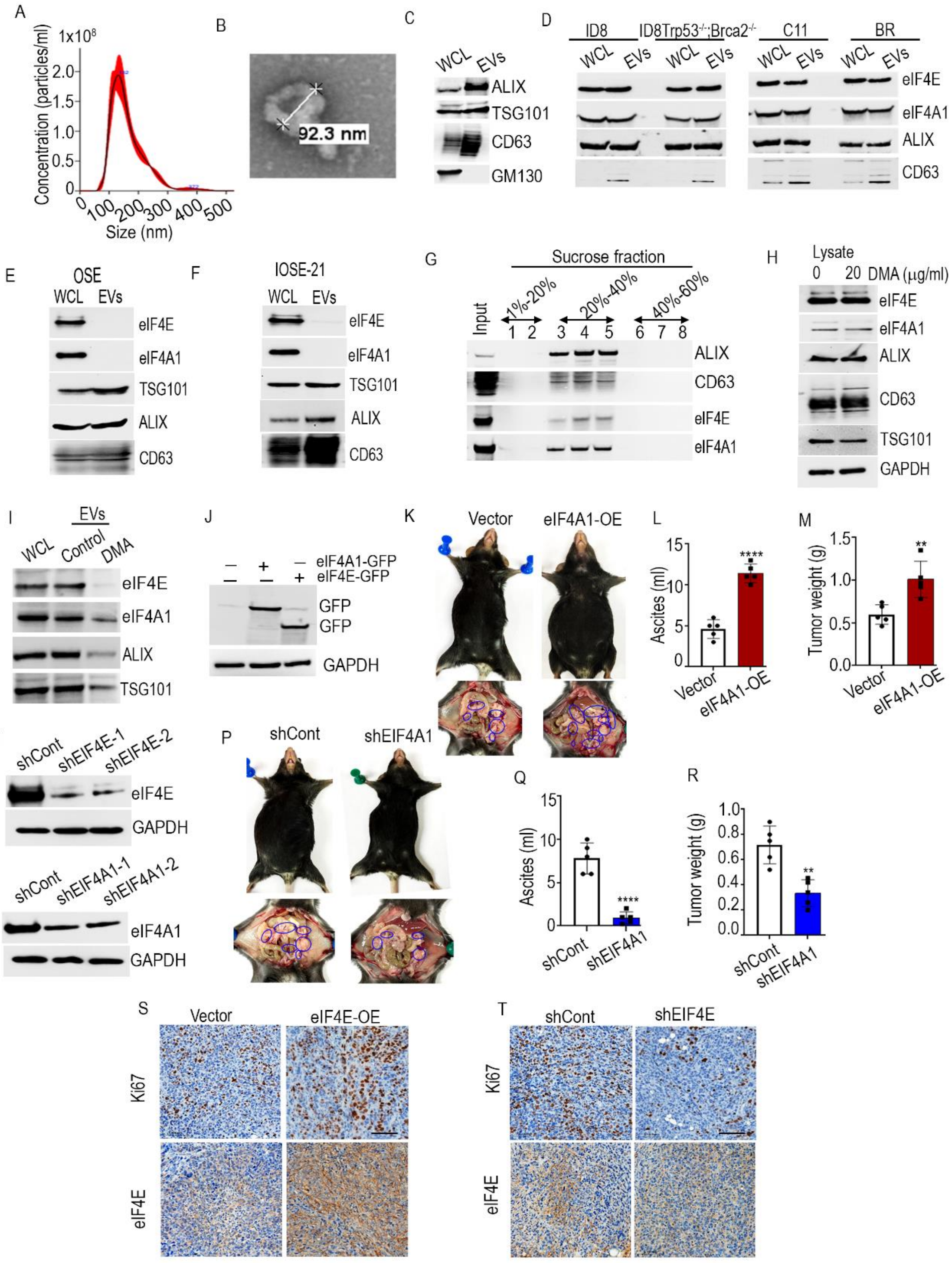

**Sup Fig-1.**

(A) Characterization of purified EVs using nanoparticle tracking (NTA).

(B) Representative TEM image of purified EVs from OVCAR5 cells.

(C) Western blotting for EVs positive (TSG101, CD63, ALIX,) and negative markers GM130.

(D-F) Western blot analysis for eIF4E and eIF4A1 proteins on whole cell lysates and isolated EVs.

(G) Western blotting using sucrose gradient fractions.

(H and I) Western blotting of DMA-treated OVCAR8 cells.

(J) Western blotting of GFP-tagged eIF4A1 and eIF4E were overexpressed in ID8Trp53$^{-/-}$;Brca2$^{-/-}$ cells

(K) Representative image of the mouse from vector control and eIF4A1 overexpressed groups. n=5.

(L) ascites volume and (M) tumor weight.

(N and O) Efficiency of eIF4E and eIF4A1 KD in ID8Trp53$^{-/-}$;Brca2$^{-/-}$ cells using shRNA (shEIF4E and shEIF4A1 respectively).

(P) Representative image of the mouse from sh-control and sh-eIF4A1 group. n=5.

(Q) ascites volume and (R) tumor weight.

(S and T) IHC analysis of tumor tissues selected from each group. Scale bar-100 µm.

The data are shown as mean ± SEM. **p<0.01,****p<0.0001 compared to control.

### Sup Fig-2a

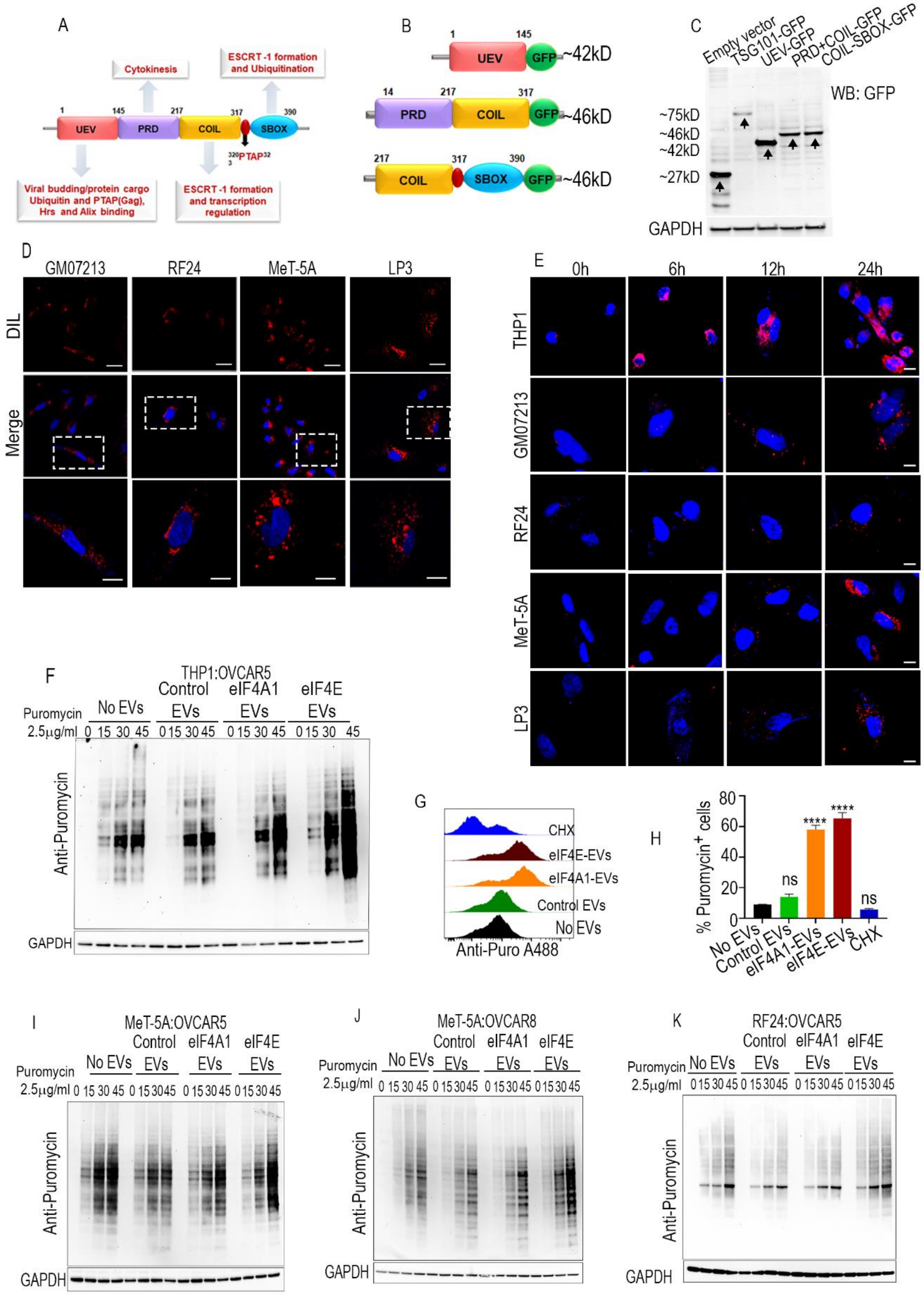

A
Cytokinesis
ESCRT-1 formation and Ubiquitination
UEV
PRD
COIL
SBOX
Viral budding/protein cargo Ubiquitin and PTAP(Gag), Hrs and Alix binding
ESCRT-1 formation and transcription regulation
B
~42kD
~46kD
~46kD
C
Empty vector
TSG101-GFP
UEV-GFP
PRD+COIL-GFP
COIL-SBOX-GFP
WB: GFP
~75kD
~46kD
~42kD
~27kD
GAPDH
D
GM07213
RF24
MeT-5A
LP3
DIL
Merge
E
0h
6h
12h
24h
THP1
GM07213
RF24
MeT-5A
LP3
F
THP1:OVCAR5
Puromycin 2.5μg/ml
No EVs
Control EVs
eIF4A1 EVs
eIF4E EVs
Anti-Puromycin
GAPDH
G
CHX
eIF4E-EVs
eIF4A1-EVs
Control EVs
No EVs
Anti-Puro A488
H
% Puromycin+ cells
ns
****
****
ns
No EVs
Control EVs
eIF4A1-EVs
eIF4E-EVs
CHX
I
MeT-5A:OVCAR5
J
MeT-5A:OVCAR8
K
RF24:OVCAR5
Anti-Puromycin
GAPDH

**Sup Fig-2b**

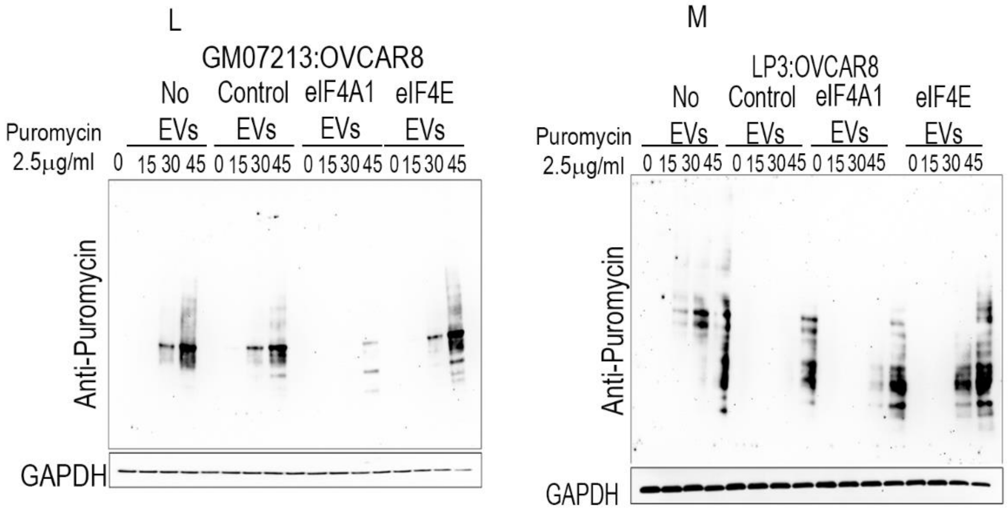


**Sup Fig-2a,b.**

(A) Schematic description of full-length TSG101.

(B) Schematic description of truncated TSG101.

(C) Western blotting showing the expression of full-length and truncated TSG101 proteins with GFP tag at C-terminal in OVCAR8 cells.

(D) The uptake of EVs by indicated cells. Scale bar 50μm and 20μm.

(E) Cells were incubated with DIL-labeled EVs for various time periods. Scale bar 50μm.

(F) SUnSET measurements of protein synthesis by western blotting in THP1-derived macrophages after treatment with EVs isolated from OVCAR5 cells.

(G and H) FACS of the puromycin-labeled THP1-derived macrophage using the anti-puromycin antibody tagged with Alexa Fluor 488.

(I-M) SUnSET measurements of protein synthesis by western blotting using indicated cells after treatment with EVs isolated from ovarian cancer cells.

Error bars indicate mean ± SEM, ****$p<0.0001$ (one-way ANOVA)

**Sup Fig-3.**

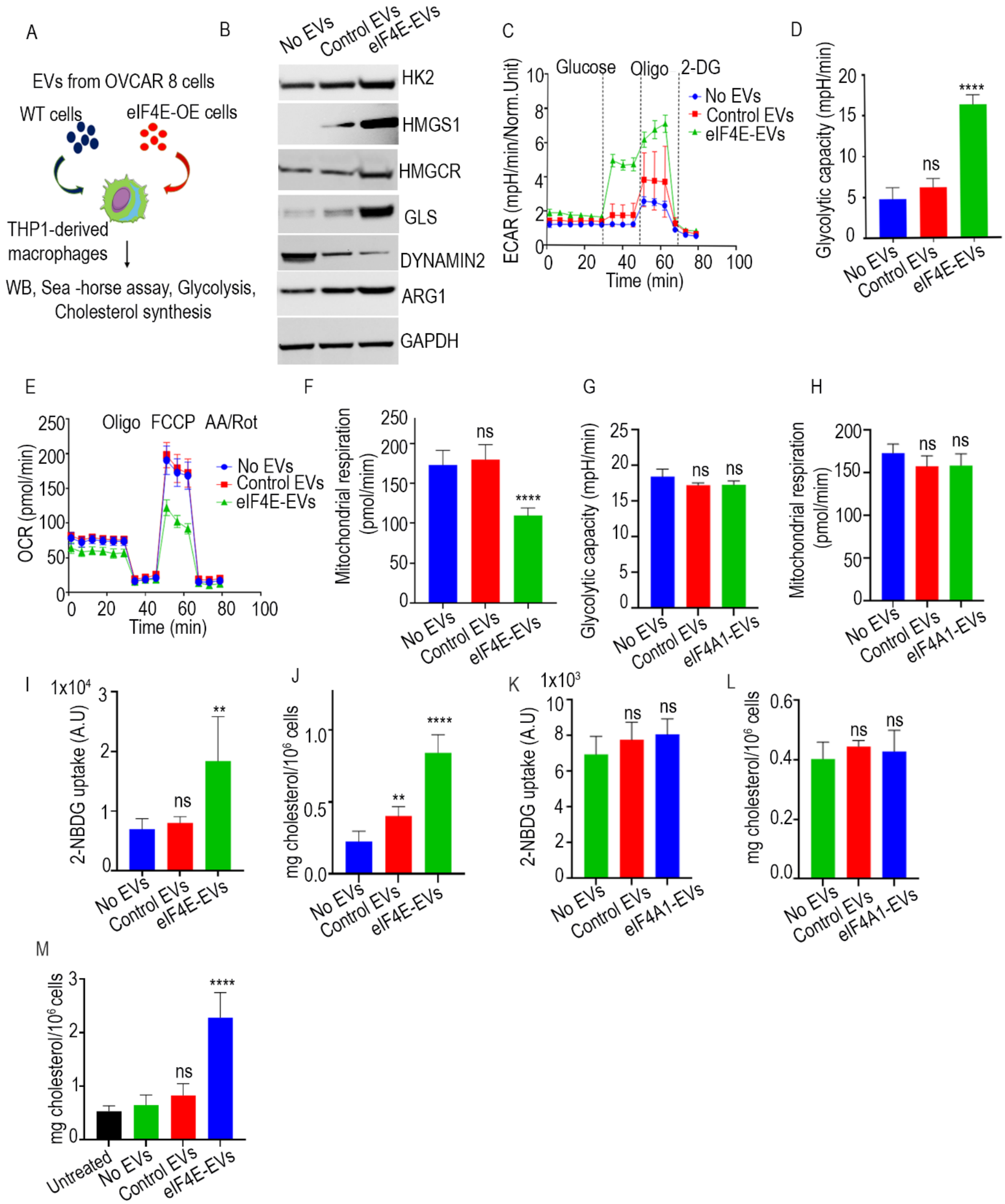
A
EVs from OVCAR 8 cells
WT cells
eIF4E-OE cells
THP1-derived macrophages
WB, Sea -horse assay, Glycolysis, Cholesterol synthesis
B
No EVs
Control EVs
eIF4E-EVs
HK2
HMGS1
HMGCR
GLS
DYNAMIN2
ARG1
GAPDH
C
Glucose
Oligo
2-DG
No EVs
Control EVs
eIF4E-EVs
ECAR (mpH/min/Norm.Unit)
Time (min)
D
Glycolytic capacity (mpH/min)
ns
****
E
Oligo
FCCP
AA/Rot
OCR (pmol/min)
Time (min)
F
Mitochondrial respiration (pmol/mim)
G
Glycolytic capacity (mpH/min)
eIF4A1-EVs
H
Mitochondrial respiration (pmol/mim)
I
1x10⁴
2-NBDG uptake (A.U)
**
J
mg cholesterol/10⁶ cells
K
1x10³
2-NBDG uptake (A.U)
L
mg cholesterol/10⁶ cells
M
mg cholesterol/10⁶ cells
Untreated

**Sup Fig-3.**

(A) Workflow representing validation of SILAC

(B) Representative western blots showing expression of HK2, HMGCS1, HMGCR, GLS, ARG-1, Dynamin-2, in macrophages treated with EVs.

(C and D) Seahorse glycolysis stress test.

(E and F) OCR measurement in eIF4E-EVs stimulated macrophages.

(G) Seahorse glycolysis stress test.

(H) OCR measurement in eIF4A1-EVs stimulated macrophages.

(I) Glucose uptake assay with 2-NBDG in macrophages stimulated with EVs.

(J) Cholesterol synthesis in macrophages by Amplex cholesterol assay.

(K) Glucose uptake assay with 2-NBDG in macrophages stimulated with EVs.

(L) Cholesterol synthesis in macrophages by Amplex cholesterol assay.

(M and N) Cholesterol measurement in OVCAR8 cells co-cultured with THP1-derived macrophages treated with EVs.

Data are representative of three independent experiments. Error bars indicate mean ± SEM, *p<0.05, **p<0.01 ****p<0.0001, (one-way ANOVA).

**Sup Fig 4**

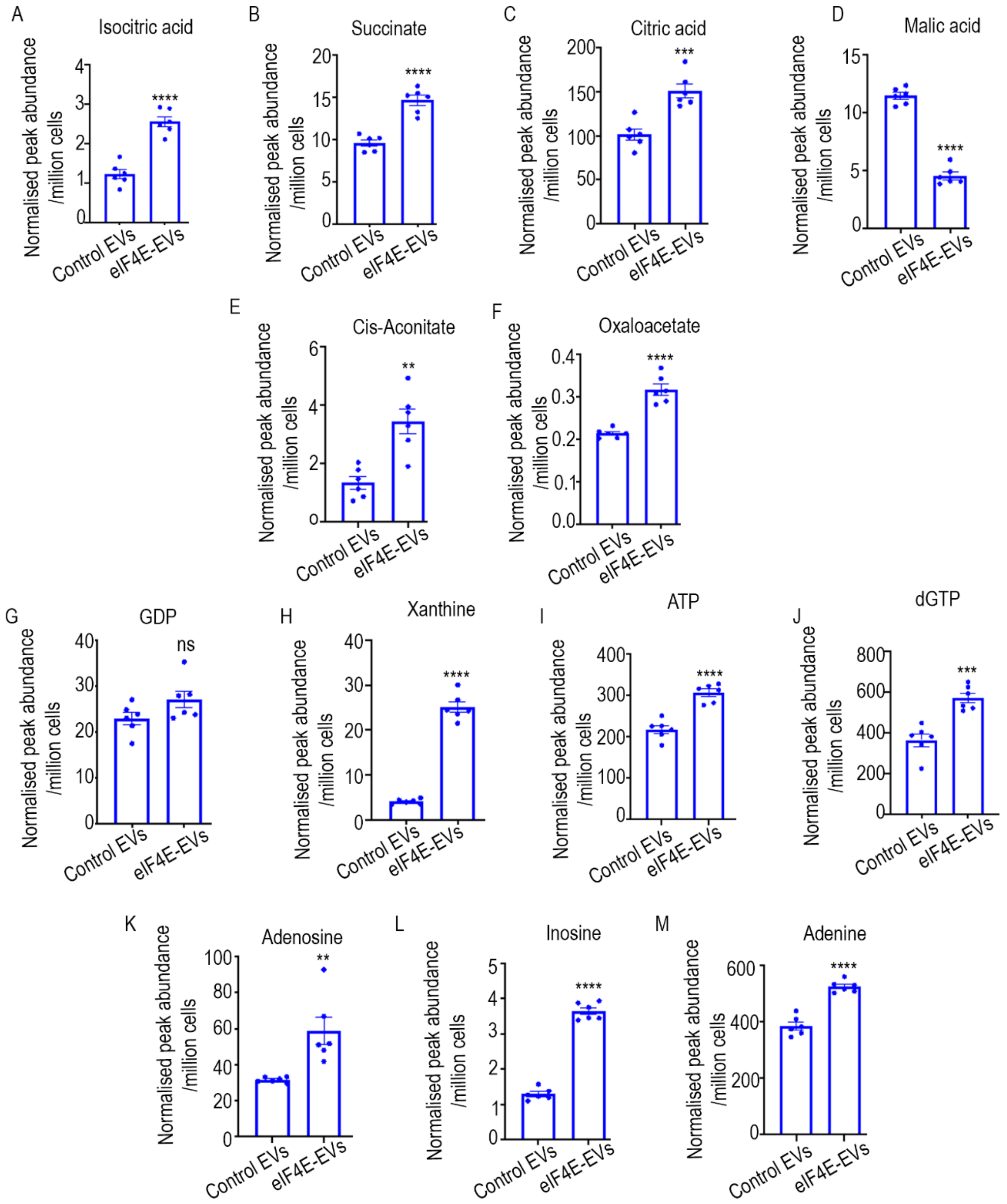


**Sup Fig 4**: (A-M) Relative metabolites levels (peak intensity normalized by internal standard and cell number in macrophages treated with control and eIF4E-EVs. (n=6). **p<0.01, ***p<0.001, ****p<0.0001, (Student's t-test). ns, non-significant.

**Sup Fig 5**

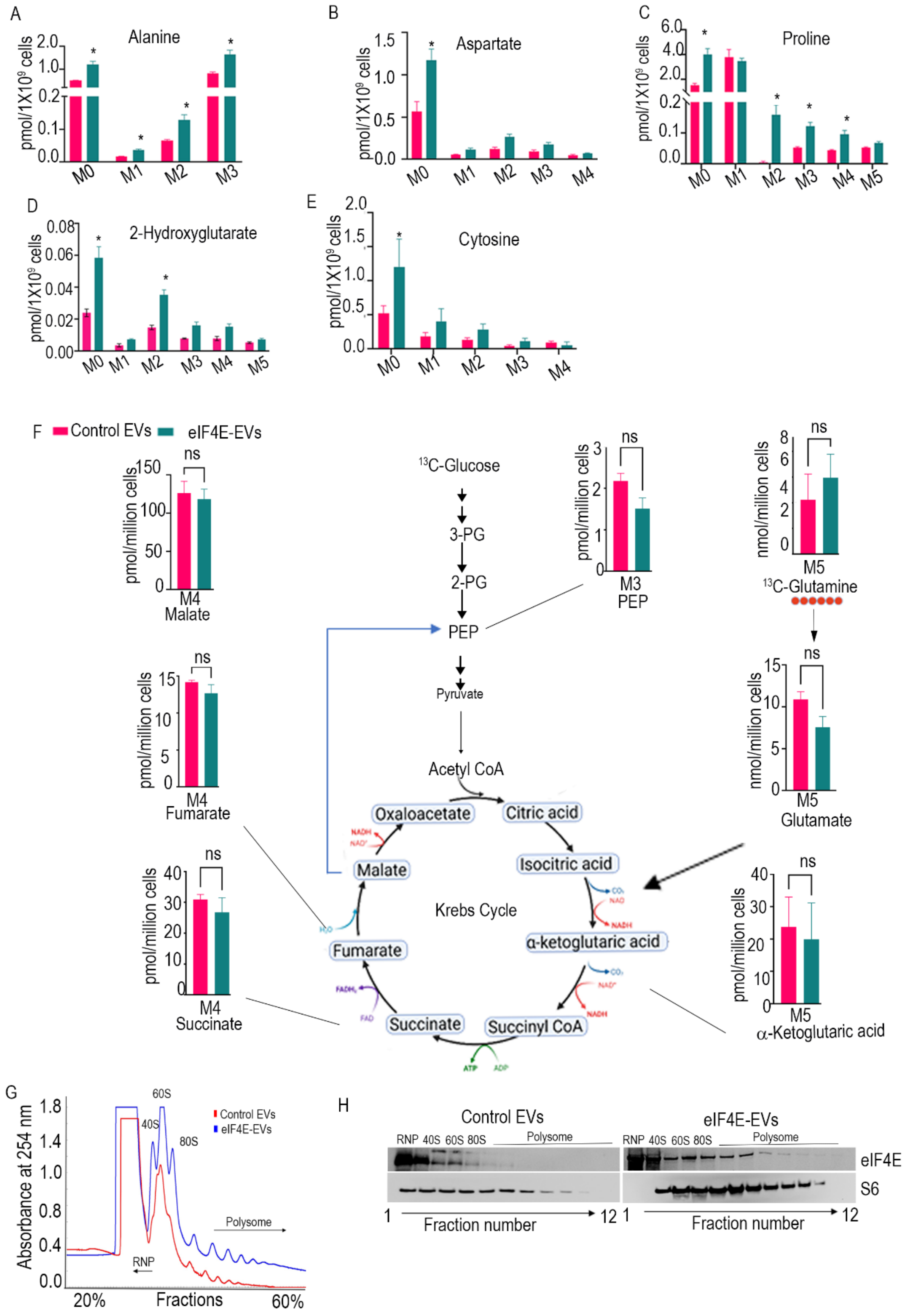

A
Alanine
pmol/1X10⁹ cells
M0 M1 M2 M3
B
Aspartate
M0 M1 M2 M3 M4
C
Proline
M0 M1 M2 M3 M4 M5
D
2-Hydroxyglutarate
M0 M1 M2 M3 M4 M5
E
Cytosine
M0 M1 M2 M3 M4
F
Control EVs
eIF4E-EVs
M4 Malate
M4 Fumarate
M4 Succinate
13C-Glucose
3-PG
2-PG
PEP
Pyruvate
Acetyl CoA
Oxaloacetate
Citric acid
Isocitric acid
α-ketoglutaric acid
Succinyl CoA
Succinate
Fumarate
Malate
Krebs Cycle
M3 PEP
M5 13C-Glutamine
M5 Glutamate
M5 α-Ketoglutaric acid
pmol/million cells
nmol/million cells
ns
G
Absorbance at 254 nm
Control EVs
eIF4E-EVs
40S
60S
80S
Polysome
RNP
Fractions
20%
60%
H
Control EVs
eIF4E-EVs
RNP 40S 60S 80S Polysome
eIF4E
S6
Fraction number

**Sup Fig 5**.

(A-E) Fractional contribution of uniformly labelled $^{13}C$ glucose in control and eIF4E-EV treated macrophages (n=3).

(F) Fractional contribution of uniformly labelled $^{13}C$ glutamine in central carbon metabolites in control and eIF4E-EV treated macrophages. n=3.

(G) Plot of the absorbance profile of fractions obtained through sucrose gradients to isolate polysomes.

(H) Western blot analysis of the protein fractions isolated from (G) was performed using the antibodies indicated.

Error bars indicate mean ± SEM, $*p<0.05$. ns, non-significant.

**Sup Fig 6**

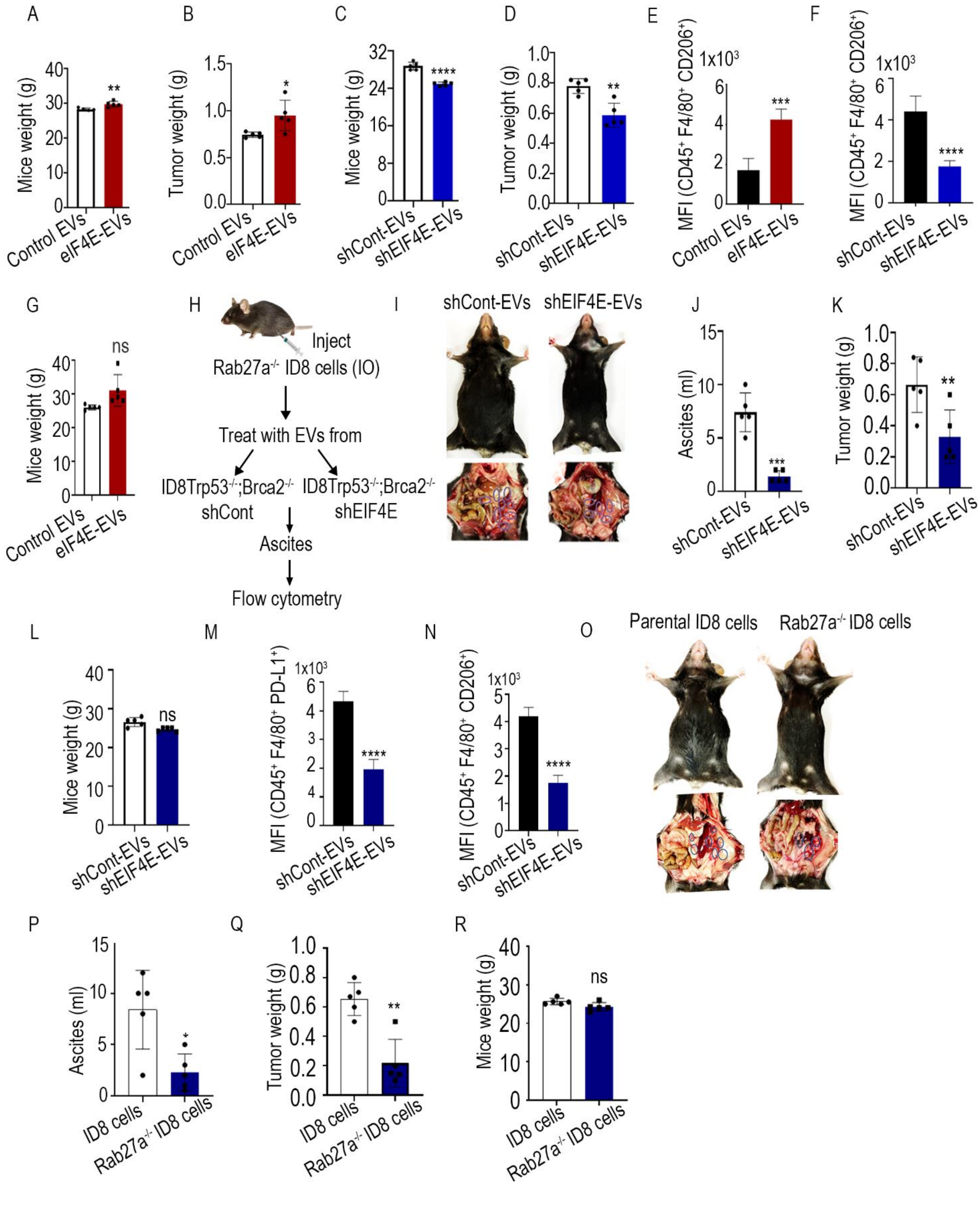

A
Mice weight (g)
Control EVs
eIF4E-EVs
B
Tumor weight (g)
C
shCont-EVs
shEIF4E-EVs
D
E
MFI (CD45+ F4/80+ CD206+)
1x10^3
F
G
ns
H
Inject
Rab27a-/- ID8 cells (IO)
Treat with EVs from
ID8Trp53-/-;Brca2-/- shCont
ID8Trp53-/-;Brca2-/- shEIF4E
Ascites
Flow cytometry
I
J
Ascites (ml)
K
L
M
MFI (CD45+ F4/80+ PD-L1+)
N
O
Parental ID8 cells
Rab27a-/- ID8 cells
P
ID8 cells
Q
R

**Sup Fig 6**

(A and C) Bar graph indicates the average mice weight.

(B and D) Average tumor weight.

(E and F) CD206 expression on macrophages in ascites samples. n=3.

(G) Bar graph indicates the average mice weight.

(H) Schematic for experimental design (I-N).

(I) Ascites accumulation in the peritoneal cavity of mice.

(J) ascites volume (K) tumor weight (L) mice weight.

(M) PD-L1 expression (N) CD206 expression on macrophages in ascites samples. n=3.

(O) Accumulation of ascites and tumor nodules in the peritoneal cavity. n=5.

(P) ascites volume (Q) tumor weight, and (R) mice weight.

Error bar represents ±, SEM, *$p<0.05$, **$p<0.01$ ***$p<0.001$ ****$p<0.0001$ (Student's t-test). ns, non-significant.

**Sup Fig 7**

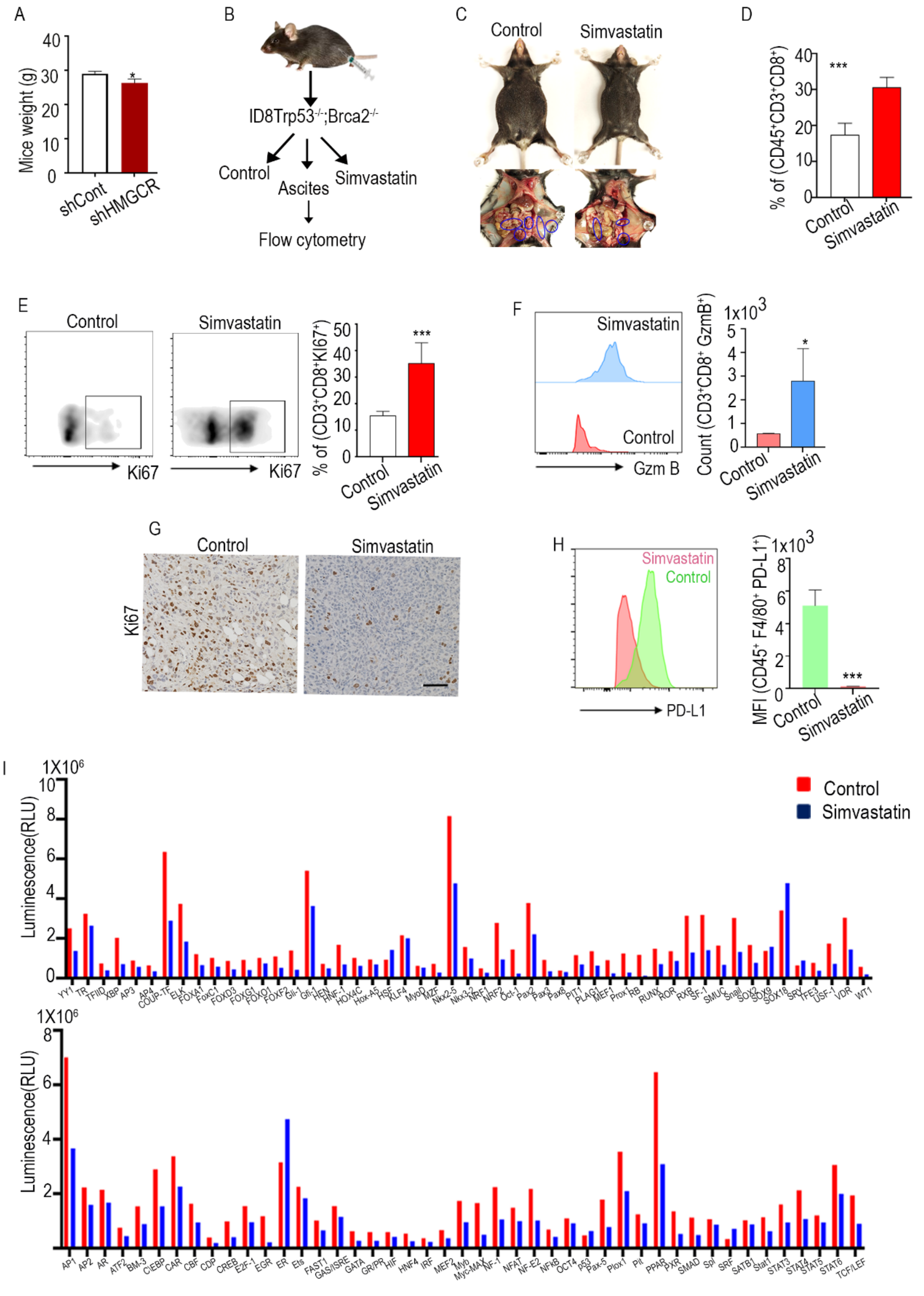

A
Mice weight (g)
shCont
shHMGCR
B
ID8Trp53-/-;Brca2-/-
Control
Ascites
Simvastatin
Flow cytometry
C
Control
Simvastatin
D
% of (CD45+CD3+CD8+)
Control
Simvastatin
E
Control
Simvastatin
Ki67
% of (CD3+CD8+KI67+)
F
Simvastatin
Control
Gzm B
Count (CD3+CD8+ GzmB+)
1x10^3
G
Control
Simvastatin
Ki67
H
Simvastatin
Control
PD-L1
MFI (CD45+ F4/80+ PD-L1+)
1x10^3
I
1X10^6
Luminescence(RLU)
Control
Simvastatin

**Sup Fig 7.**

(A) Bar graphs indicate mice weight.

(B) Schematic of the simvastatin treatment (10mg/kg)

(C) Peritoneal cavity of mice showing ascites accumulation and tumor nodules.

(D) Flow cytometric quantification of $CD8^+$ cells among $CD45^+$ and $CD3^+$ cells in ascites.

(E and F) Quantification of $Ki67^+$ cells and $GzmB^+$ cells among $CD8^+$ T cells in ascites.

(G) IHC analysis of tumor tissues

(H Flow cytometry analysis of PD-L1 expression in peritoneal macrophages.

(I) Bar graph gives the results of a transcription factor array.

Error bar represents ±, SEM, *$p<0.05$, ***$p<0.001$ (Student's t-test)

**Sup Fig 8**

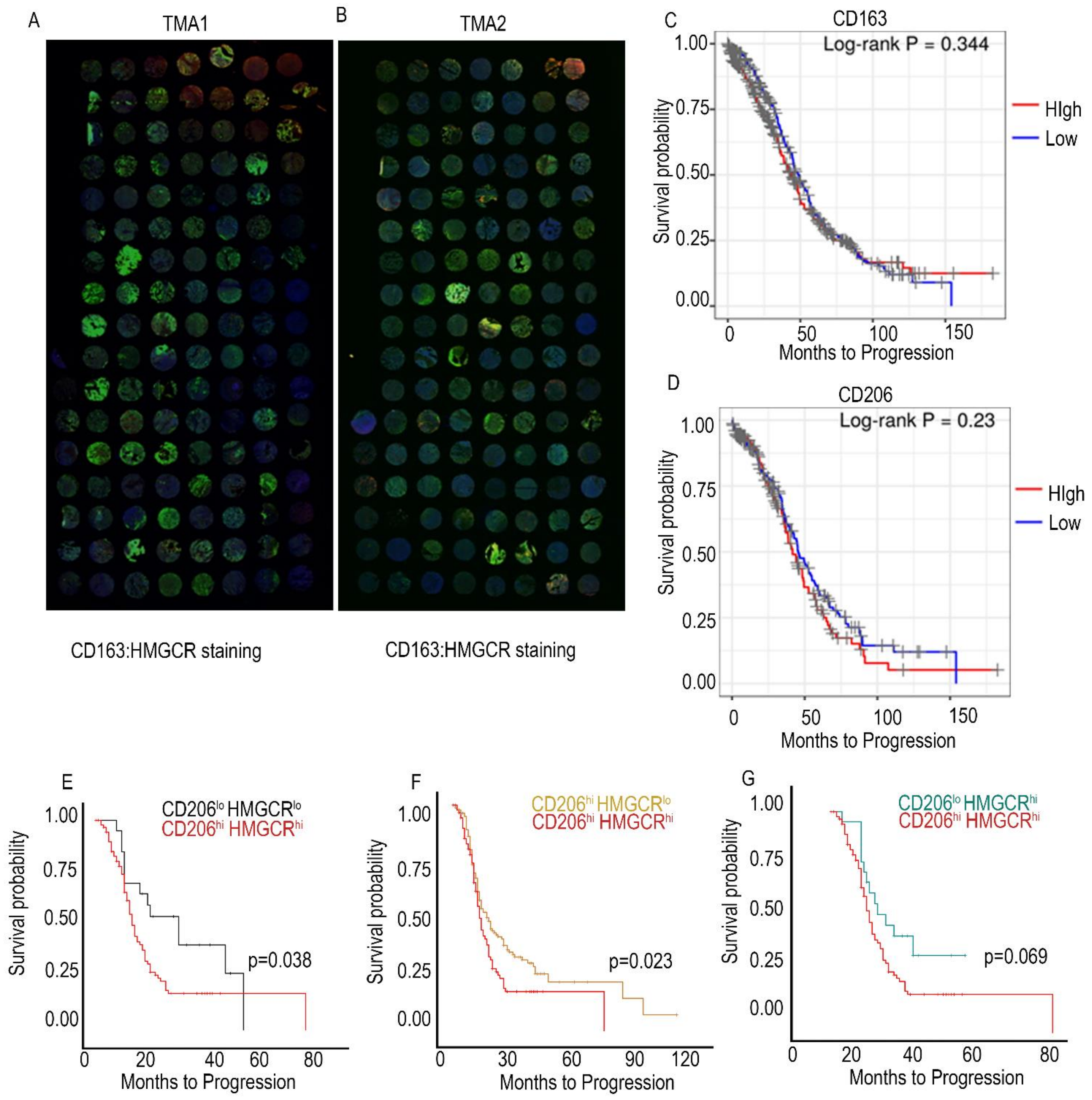
A
TMA1
B
TMA2
CD163:HMGCR staining
CD163:HMGCR staining
C
CD163
Log-rank P = 0.344
HIgh
Low
Survival probability
Months to Progression
D
CD206
Log-rank P = 0.23
HIgh
Low
Survival probability
Months to Progression
E
CD206lo HMGCRlo
CD206hi HMGCRhi
p=0.038
Survival probability
Months to Progression
F
CD206hi HMGCRlo
CD206hi HMGCRhi
p=0.023
Survival probability
Months to Progression
G
CD206lo HMGCRhi
CD206hi HMGCRhi
p=0.069
Survival probability
Months to Progression

**Sup Fig 8.**

(A and B) Co-Immunofluorescence staining for HMGCR and CD163 in the whole slide of human ovarian cancer tissue microarray.

(C) Survival curves of high CD163 vs low CD163 expression patients.

(D) Survival curves of high CD206 vs low CD206 expression patients.

(E) Survival curves of high HMGCR and high CD206 expression vs. low HMGCR and low CD206 expression.

(F) Survival curves of high HMGCR and high CD206 expression vs. low HMGCR and high CD206 expression

(G) Survival curves of high HMGCR and high CD163 expression vs. high HMGCR and low CD206 expression
